# *In vivo* mapping of the deep and superficial white matter connectivity in the chimpanzee brain

**DOI:** 10.1101/2023.08.25.554775

**Authors:** Maëlig Chauvel, Ivy Uszynski, Bastien Herlin, Alexandros Popov, Yann Leprince, Jean-François Mangin, William D. Hopkins, Cyril Poupon

**Author notes:** Maëlig Chauvel; Cyril Poupon.

## Abstract

Mapping the chimpanzee brain connectome, and comparing it to humans, is key to our understanding of similarities and differences in primate evolution that occurred after the split from their common ancestor around 6 million years ago. In contrast with macaque species brain studies, less studies have specifically addressed the structural connectivity of the chimpanzee brain and its comparison with the human brain. Most comparative studies in the literature focus on the anatomy of the cortex and deep nuclei to evaluate how their morphometry and asymmetry differs from that of the human brain, and some studies have emerged concerning the study of brain connectivity between primates. In this work, we established a new white matter atlas of the deep and superficial white matter structural connectivity in chimpanzees. In vivo anatomical and diffusion weighted magnetic resonance imaging (MRI) data were collected on a 3 Tesla magnetic resonance imaging (MRI) system in 39 chimpanzees. These datasets were subsequently processed using a dedicated fiber clustering pipeline adapted to the chimpanzee brain enabling us to create two novel deep and superficial white matter connectivity atlases representative of the chimpanzee brain. These atlases provide the scientific community with an important and novel set of reference data for understanding the commonalities and differences of the structural connectivity between the human and chimpanzee brains, which will contribute to a better understanding of the hominin brain evolution.

## 1 Introduction

Humans and chimpanzees split from a common ancestor between 6 to 8 million years ago (Amster and Sella 2016) and share as much as 96% of the same genome sequence(Suntsova and Buzdin 2020, Consortium et al. 2005). Compared to more distantly related primate species, chimpanzees and humans also show a number of anatomical, behavioral and cognitive similarities that were likely manifest in their common ancestor such as complex manual manipulative skills, tool use, and complex non-verbal gestural communication (Schwartz and Beran 2022, Sherwood et al. 2012,Shumaker, Walkup, and Beck 2011). Despite these similarities, it is also clear that humans evolved unique anatomical, neurological, cognitive and motor abilities since their split from the common ancestor of chimpanzees: these abilities include but are not limited to bipedalism, complex vocal communication comprising language and speech, and higher order cognitive and motor functions (Bradshaw and Rogers 1992, Fitch 2010, Tomasello and Carpenter 2007, Arbib, Liebal, and Pika 2008, Rizzolatti and Arbib 1998).

What neurological changes occurred during hominin evolution that accounts for the emergence of uniquely human abilities (and disorders) has been a central focus of comparative neuroscience research. Historically, comparative neuroscience research has focused on either (1) allometric changes in whole brain or region specific volumes between different primate species (Hopkins 2007,James K Rilling and Insel 1999a, James K Rilling and Insel 1999b, Semendeferi et al. 2001,Semendeferi et al. 1998) or (2) microscopic and cellular changes between primate species for different brain regions (Schenker et al. 2008, Sporns, Tononi, and Kötter 2005). With the emergence of *in vivo* MRI brain imaging, comparative neuroscience research has accelerated and expanded to include larger sample sizes within different species and to include more diverse measures of neural organization.

Diffusion MRI is now a well-established MRI tool to explore the brain microstructure through the observation of the anisotropy of Brownian motion of water molecules present in tissues. To date, it remains the only method for in vivo imaging of brain anatomical connectivity. Sulcal-gyral anatomy and structural connectivity are intimately linked, with brain functions resulting from the connection of cortical regions and/or deep gray matter regions via axonal fiber bundles populating the cerebral white matter, thus establishing functionally and anatomically connected networks. One of the great challenges of comparative neuroscience from an evolutionary point of view, is to build and compare models of the brain connectome in order to be able to compare species. The neuroimaging community has been working actively for almost twenty years to establish the human brain structural connectome. Several atlases of long fascicle anatomical connectivity have been proposed following different methodological approaches (Catani and DeSchotten 2008, P. Guevara et al. 2012, Yendiki et al. 2011). More recently, new atlases of the superficial anatomical connectivity with more challenging inference have been proposed for humans that allow for a finer exploration of the link between sulcalogyral anatomy and subcortical connectivity (M. Guevara et al. 2020). These methodological advances in studies with humans have opened new ways for examining the connectome of other animal species, including chimpanzees. In chimpanzees, a first anatomically-informed atlas was recently released by Bryant, et al. (K. Bryant et al. 2020), from a cohort of 29 female individuals scanned using diffusion MRI. This atlas was built from anatomically-informed regions of interest corresponding to long fascicle crossing points in each individual and allowed for the identification of 24 fascicles. To our knowledge, this or any other atlas for chimpanzees does not include any information about the superficial structural connectivity. Furthermore, the chosen cohort was exclusively composed of females which can be a source of bias with respect to inter-sex variation. Indeed, previous work have pointed out inter-sex differences in the chimpanzee brain morphology, whether concerning the planum parietale and temporale or the corpus callosum (Taglialatela, Dadda, and Hopkins 2007, Hopkins et al. 2016).

In this work, we propose a refined long white matter bundles atlas and establish for the first time an atlas of the superficial white matter atlas of the chimpanzee brain. Having access to both long and superficial white matter connectivity will more thoroughly contribute to the identification of similarities and differences in connectivity between humans and chimpanzees. The construction of these novel atlases relies on the development of a dedicated fiber clustering pipeline which does require only a very few anatomical priors, and thus, allows to automatically infer the superficial connectivity of the chimpanzee brain whose anatomy is largely unknown today, as for humans, contrarily to the well known deep structural connectivity from which anatomical landmarks are well identified.

## 2 Materials and Methods

### 2.1 Data Collection

We considered data coming from 39 healthy in vivo chimpanzees including 23 females and 16 males, imaged between 9 and 35 years old (mean = 19 years old) and housed at the Emory National Primate Research Center/ENPRC. Chimpanzee MRI scans were obtained from a data archive of scans acquired prior to the 2015 implementation of US Fish and Wildlife Service and National Institutes of Health regulations governing research with chimpanzees. All the scans reported in this publication were completed by the end of 2012 and have been used in previous studies (e.g. Vickery et al. 2020 or Bogart et al. 2012). Each individual was scanned using a 3 Tesla Trio MRI system (Siemens, Erlangen) using a birdcage coil with a dedicated imaging protocol including : a high resolution 3D T1-weighted inversion recovery fast gradient echo anatomical scan (MPRAGE sequence) of 0.625mm isotropic resolution (echo time TE/TR=4.38ms/2600ms, inversion time TI=900ms, flip angle FA=8°, matrix size 256 x 256, read bandwidth RBW=130Hz/pixel), and five diffusion-weighted MRI scans using a 2D single-shot twice refocused Pulse gradient spin echo (PGSE) echo-planar (EPI) sequence of 1.9mm isotropic resolution (TE/TR=86ms/6s, flip angle FA=90°, read bandwidth RBW=1563Hz/pixel, matrix size 128×128, FOV=243.2mm) at b=1000s/mm² along 60 diffusion directions. All procedures were carried out in accordance with protocols approved by YNPRC and the Emory University Institutional Animal Care and Use Committee.

### 2.2 Data preprocessing, local modeling and fiber tractography

Anatomical and diffusion data were processed using a Python pipeline dedicated to the chimpanzee species using the in-house C++ Ginkgo toolbox available at https://framagit.org/cpoupon/gkg. All T1-weighted images of the 39 chimpanzees’ brains were matched to the Juna.chimp chimpanzee template (template release : https://www.chimpanzeebrain.org/) Vickery et al. 2020). To this aim, direct and inverse non-linear 3D registration transformations from each individual brain T1-weighted scan to the Juna.chimp template were computed using the ANTs software (Avants, Tustison, and Song 2008), relying on a Symmetric Normalization (SyN) with a mutual information similarity criteria. The Juna.Chimp template is released with a high spatial resolution suitable for our project and corresponds to the size of a standard chimpanzee brain (from anterior to posterior =106mm; from up to bottom = 72mm; from left to right = 87mm). It also includes a cortical parcellation of interest to establish the superficial white matter atlas. At the individual scale, diffusion-weighted MRI images were first corrected for susceptibility artifacts with the Top-Up method described in Andersson, Skare, and Ashburner 2003, and implemented in FSL (Smith et al. 2004). The corrected reference dMRI volume acquired at b=0s/mm2 was matched to the anatomical T1-weighted MRI using a rigid transformation computed with the Ginkgo registration tool relying on a mutual information criterion. Individual maps of local orientation distribution functions (ODF) were reconstructed from the corrected dMRI scans using the analytical Q-ball model (Descoteaux, Angelino, et al. 2007) at spherical harmonics order SH=6 with a Laplace-Beltrami regularization factor λ = 0.006. A whole-brain streamline regularized deterministic tractography algorithm (1 seed/voxel, forward step 0.4mm, aperture angle 30°, lower GFA = 0.15)(Perrin et al. 2005) was then applied to each ODF map to generate streamlines within a mask corresponding to the brain established from the anatomical MRI, yielding the 39 chimpanzees’ individual tractograms. Streamlines whose length did not belong to the 7mm-133 mm range were filtered out.

### 2.3 Two-fold fiber clustering

Because fiber tracking methods can produce millions of streamlines per subject, establishing white matter bundle atlases from a group of subjects is a difficult task from a computational point of view due to the large size of the connectivity matrices involved at the group level, even if the group size remains small. The approach that was originally proposed in Guevara et al. 2012 (P. Guevara et al. 2012) and that we also used is to decompose the problem into several steps in order to take benefit from different strategies to subsequently reduce the dimension of the problem. This strategy is based on a first clustering step, called intra-subject fiber clustering, establishing fascicles (e.g. individual fiber clusters) at the subject level, followed by a second step, called inter-subject fiber clustering” providing fascicle clusters at the population level.

#### 2.3.1 Intra-subject fiber clustering

The intra-subject fiber clustering relies on 8 consecutive steps (see figure 1):

- step 1: streamlines reconstructed using fiber tracking from each individual diffusion MRI dataset are transformed into the Davi130 template space using a diffeomorphic transformation calculated with the ANTs toolbox (Avants, Tustison, and Song 2008);
- step 2: transformed streamlines are separated into four subgroups corresponding to the brain area they mostly belong to, eg the right or left hemisphere, the inter-hemispheric region or the brainstem/cerebellum region; this step is achieved using a mask of these regions defined from the Davi130 atlas (Vickery et al. 2020);
- step 3: each streamline subgroup is then subdivided into nine new subgroups corresponding to 9 equally spaced fiber length ranges between 7mm and 133mm;
- step 4: for each of the 9 subgroups and each brain area, a fiber density mask is computed and thresholded to provide a binary mask corresponding to the brain region populated by the streamline of the chosen length range;
- step 5: a k-means algorithm is used to compute a parcellation of each binary mask with parcels sharing of similar size;
- step 6: an affinity matrix is computed between parcels that simply quantify the level of connectivity of each pair of parcels;
- step 7: a hierarchical clustering algorithm is applied to this affinity matrix providing a dendrogram allowing to define clusters of strongly connected parcels;
- step 8: parcel clusters are finally intersected with the input streamline subgroup of interest, yielding the set of fiber clusters (or fascicles) corresponding to the fiber length range and brain area of interest, and the centroid of each fascicle corresponding to the streamline of the fascicle being the closest to all the other streamlines (using a symmetric mean of mean closest point distance) is computed to provide a synthetic representation of the fascicle that will be used during the next stage operated at the population level.

**Figure 1:**
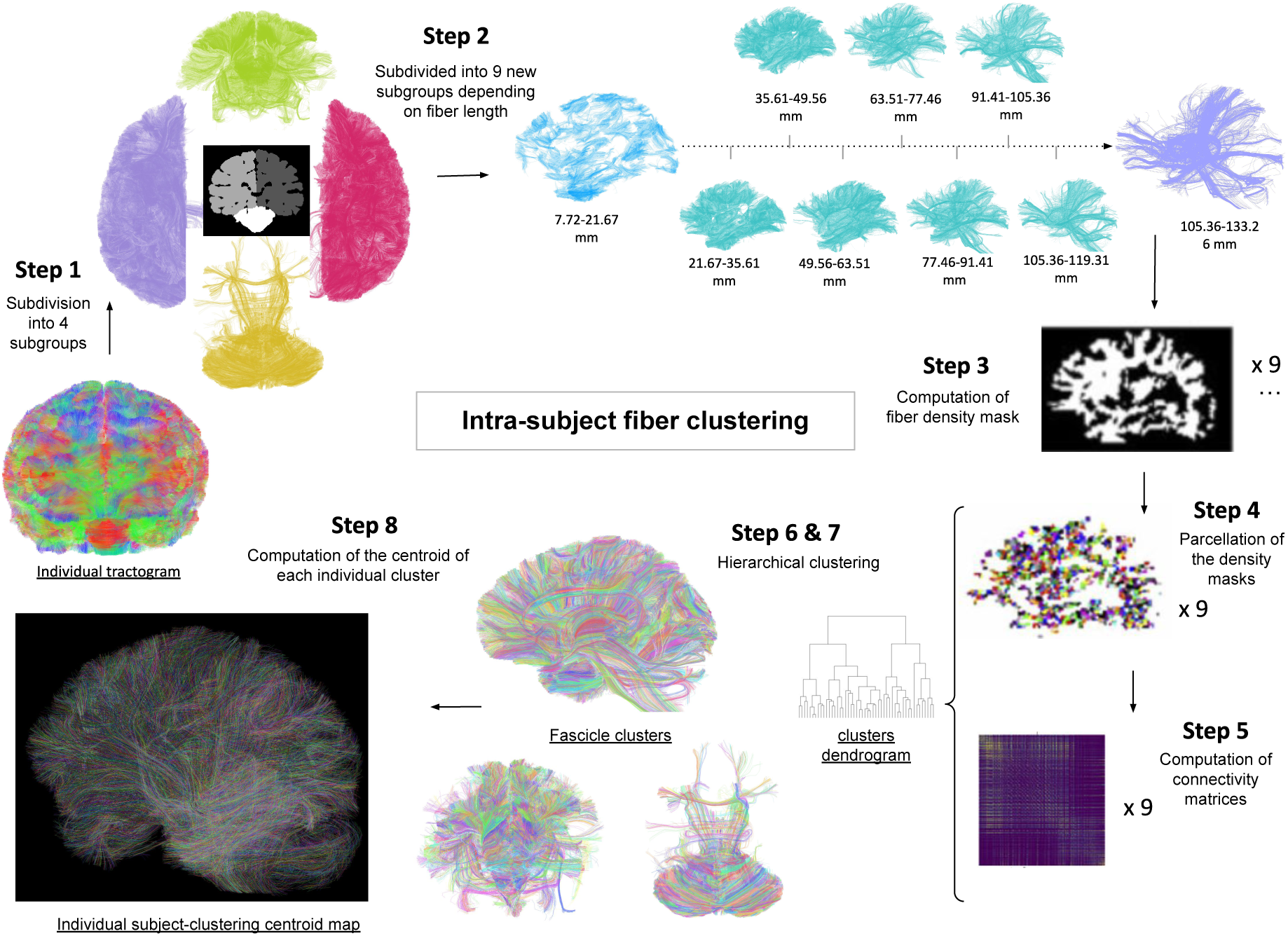
Intra-subject fiber clustering pipeline

#### 2.3.2 Inter-subject fiber clustering

The inter-subject relies on 5 consecutive steps (see figure 2):

- step 1: the centroids of all subjects collected during the first stage and providing a synthetic representation of all the individual fascicles are resampled to further reduce their representation to 21 equally spaced points, thus allowing to drastically speed-up the computation of the distance between two centroids;
- step 2: all 21-point resampled centroids representing the individual fascicles are aggregated into 4 centroid maps corresponding to the left hemisphere, the right hemisphere, the inter-hemispheric region and the brainstem/cerebellum region;
- step 3: an affinity matrix is computed for each of these 4 centroid maps using a corrected pairwise maximum Euclidean distance between centroids. The objective of the correction is to take into account the actual morphometric difference between “long” and “short” white matter bundles: long bundles can have different extremity decussing while short bundles show less dispersed connections at their extremities. The implemented distance correction and parameters optimization are further detailed in the *Supplementary information* section. Using this modified distance, an affinity measure can be easily computed for all pairs of centroids using a Gaussian kernel 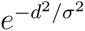 (with affinity variance equal to *σ*^2^ = 140625*mm*^2^). The affinity of a centroid pair is stored in a sparse matrix when its value is greater than a specific threshold equal to 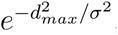, *d_max_* rep-resenting the maximum distance allowed to aggregate centroids, and considered as null otherwise;
- step 4: a density-based spatial clustering (DBSCAN)(Ester et al. 1996) of each of the 4 centroid affinity matrices is subsequently performed providing 4 sets of fascicle clusters for each brain region including the left hemisphere, the right hemisphere, the interhemispheric region and the brainstem/cerebellum region. This step yields to 4 subsets of streamlines: 2 subsets corresponding to the streamlines belonging exclusively to the two hemispheres respectively, 1 subset corresponding to the cerebellum and brainstem, and one last subset corresponding to the streamlines spreading over the 2 hemispheres, which mainly correspond to the commissural fibers and anterior/posterior commissures;
- step 5: Resulting fascicle clusters are discarded if they do not represent at least 40 percent of the subjects (percentage chosen considering the intra-subject variability). Discarded centroids are reintegrated to the kept fascicle clusters if their distance to the closest cluster remained below a maximum distance of 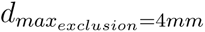.

**Figure 2:**
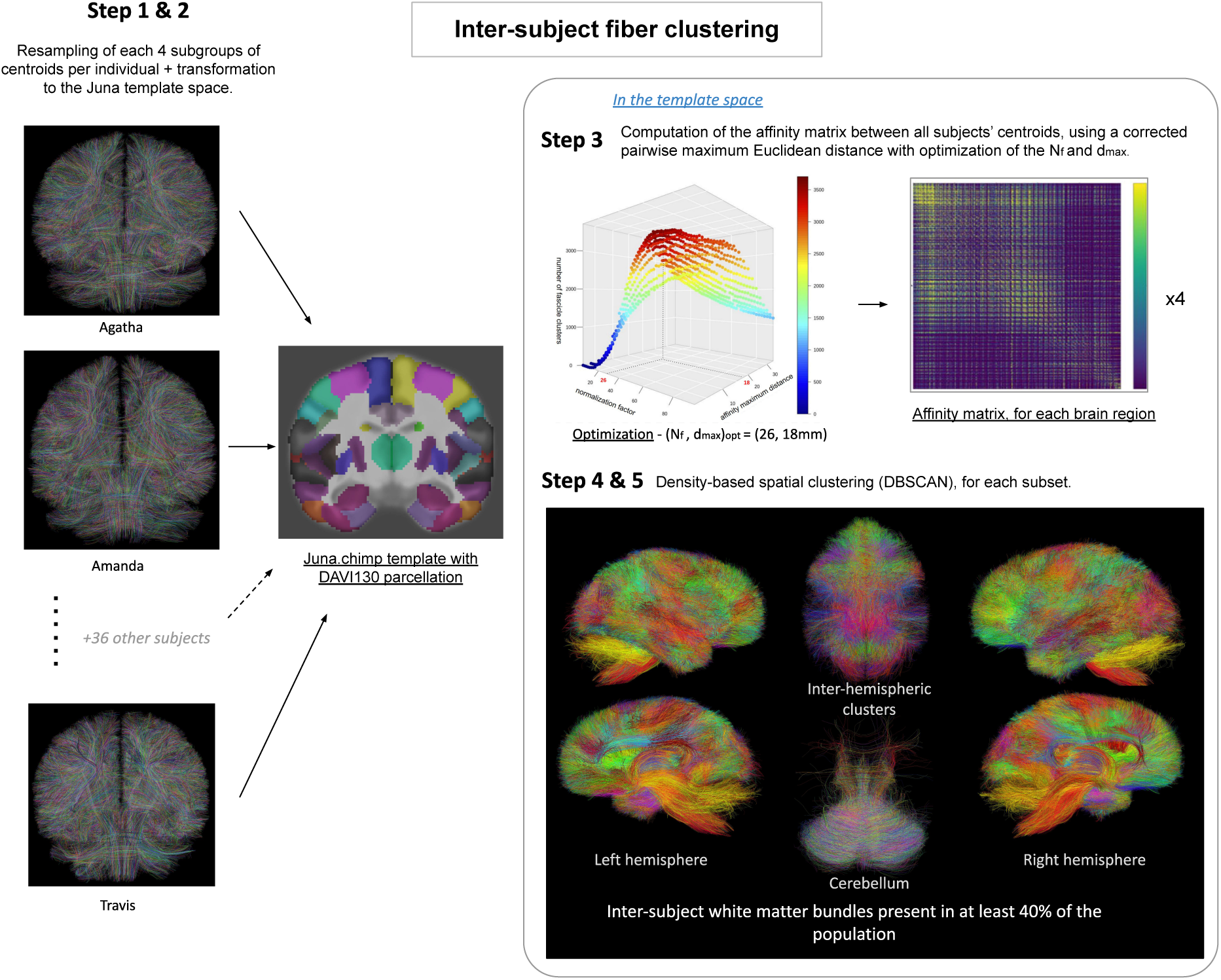
Inter-subject fiber clustering pipeline

### 2.4 Atlasing

Fascicle clusters stemming from the two-step clustering stage were separated into long or superficial fascicle clusters according to their average length with respect to an empirically chosen threshold of 50mm established from the histogram of fiber lengths at the population scale. Fascicle clusters whose lengths were under this threshold were considered ‘superficial’ whereas those whose lengths were above this threshold were considered ‘deep’.

#### 2.4.1 Design of the deep white matter bundle atlas

Deep white matter fiber bundles were designed from the aggregation of the fascicle clusters obtained at the group level, whose lengths range from 50 millimeters to 133 millimeters. The longest fascicle clusters stemming from the cross-subject clustering step were easy to select to form the chimpanzee long white matter atlas. The longest fascicles did not require the definition of regions of interest to be selected, they were already corresponding to fascicle clusters and were therefore labelled after the visual inspection by two independent neuro-anatomists (BH and WH).

#### 2.4.2 Design of the superficial white matter bundle atlas

After manually discarding fascicle clusters showing artifactual trajectories fascicle clusters were further merged using a distance criterion, in order to obtain final sets of superficial fascicle clusters relevant from an anatomical point of view. Superficial clusters were then selected using 76 cortical regions from the Davi130 parcellation atlas as can be seen in figure 3 specifically defined for the chimpanzee brain (Vickery et al. 2020). Each superficial cluster connecting two parcels A and B was attributed a name following the syntax A_B_Id (Id corresponding to the cluster index in the set of all clusters connecting parcels A and B).

**Figure 3:**
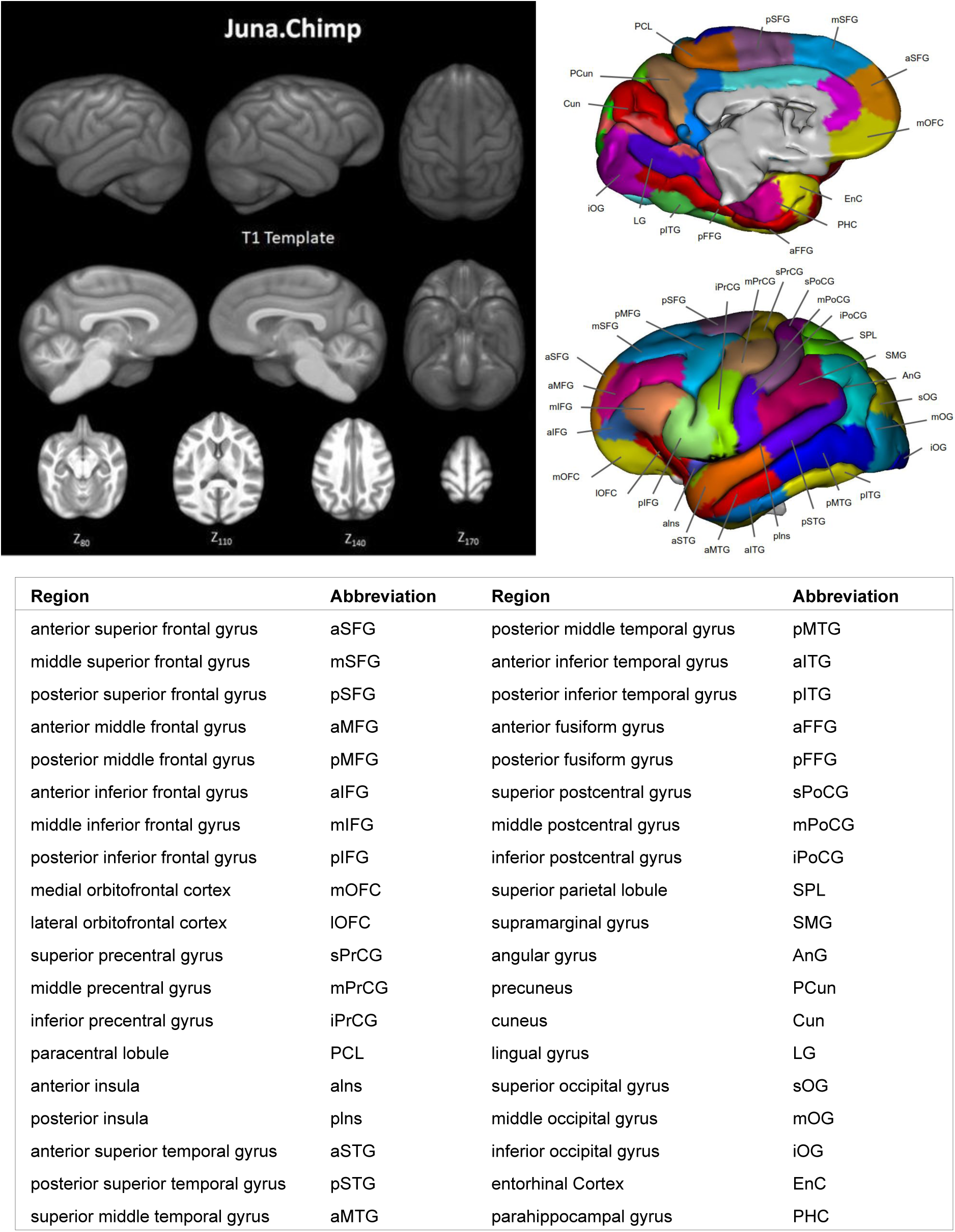
The Juna.chimp template and selected cortical regions. (Top, left): axial, coronal and sagital renderings of the template, (Top, right): the 38 cortical regions selected on each hemisphere from the DAVI130 labelling. (Bottom): table of the cortical regions selected and corresponding abbreviations. Adapted from Vickery et al. 2020.

## 3 Results

The first clustering stage resulted in fascicle sets composed of 19,9201,273 individual fibers including 10,0161,273 individual fibers in the left hemisphere and 9,9041,169 in the right hemisphere, showing a slight asymmetry in favor of the left hemisphere. The inter-subject fiber clustering yielded a total of 4862 fascicle clusters present in at least 40% of the population, subdivided into 2420 fascicle clusters for the left hemisphere and 2438 fascicle clusters for the right hemisphere (see figure 4,a.).

**Figure 4:**
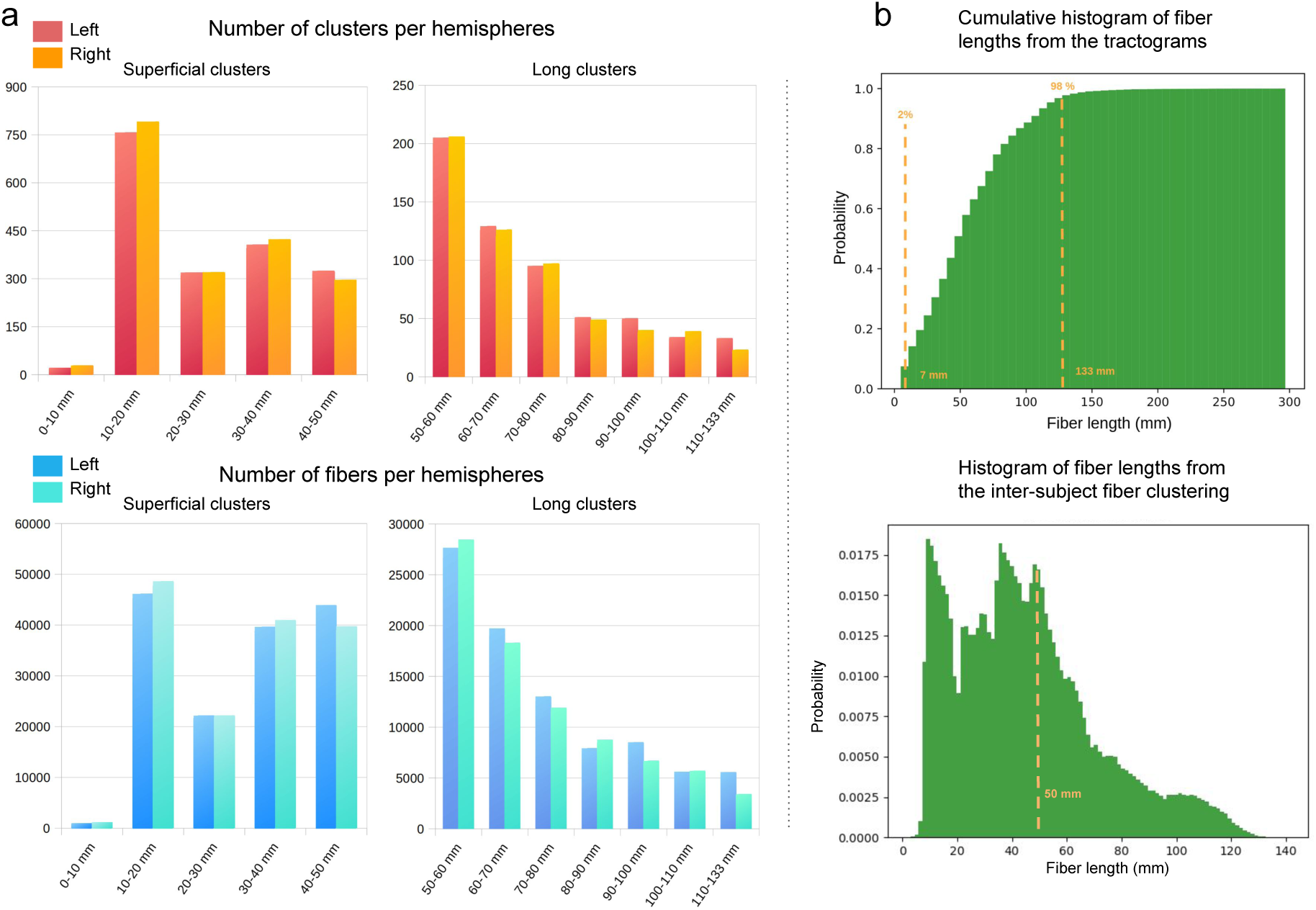
Histograms of the number of clusters relatively to the fibers length. a) Histograms of the number of clusters and fibers relatively to the fibers length, b) Cumulative histogram of fiber lengths from the tractograms allowing to select fibers from 7mm to 133mm (top) and histogram of fiber lengths from the inter-subject fiber clustering allowing partially the discrimination between deep and superficial fibers (bottom)

### 3.1 deep white matter bundle atlas of the chimpanzee brain

“Deep” bundles were defined as (or comprised of) clusters with fascicle lengths that exceeded 50 mm (see Figure 4.b). This threshold resulted from a trade-off between a visual inspection and a brain size-based scaling of the length threshold conventionally used in the literature for the human brain typically between 80 and 90 mm (Labra Avila 2020) to separate long-associative white matter bundles from superficial white matter bundles. Using this criteria, four classes of bundles comprised of 46 long white matter bundles were identified (see figure 5, and Supplementary Information SI-3 for detailed images of each tract.) in the chimpanzees including: the inferior fronto-occipital fasciculus, the middle longitudinal fasciculus, the arcuate fasciculus, the frontal aslant, the uncinate fasciculus, the fornix, the cingulum (dorsal and ventral), the motor and somato-sensory ascending and descending fibers (cortico-spinal tract and thalamo-cortical radiations), the corpus callosum, the anterior and posterior commissures, the ventral visual stream, and the optic radiations. In order to remove any specificity related to putative lateralization effects (such as for the corticospinal tract), the established white matter bundle atlas was further symmetrized with respect to the two hemispheres to avoid any bias related to the dominant laterality of the subjects used to create the atlas. The four classes of bundles as they appear in our data are described below.

**Figure 5:**
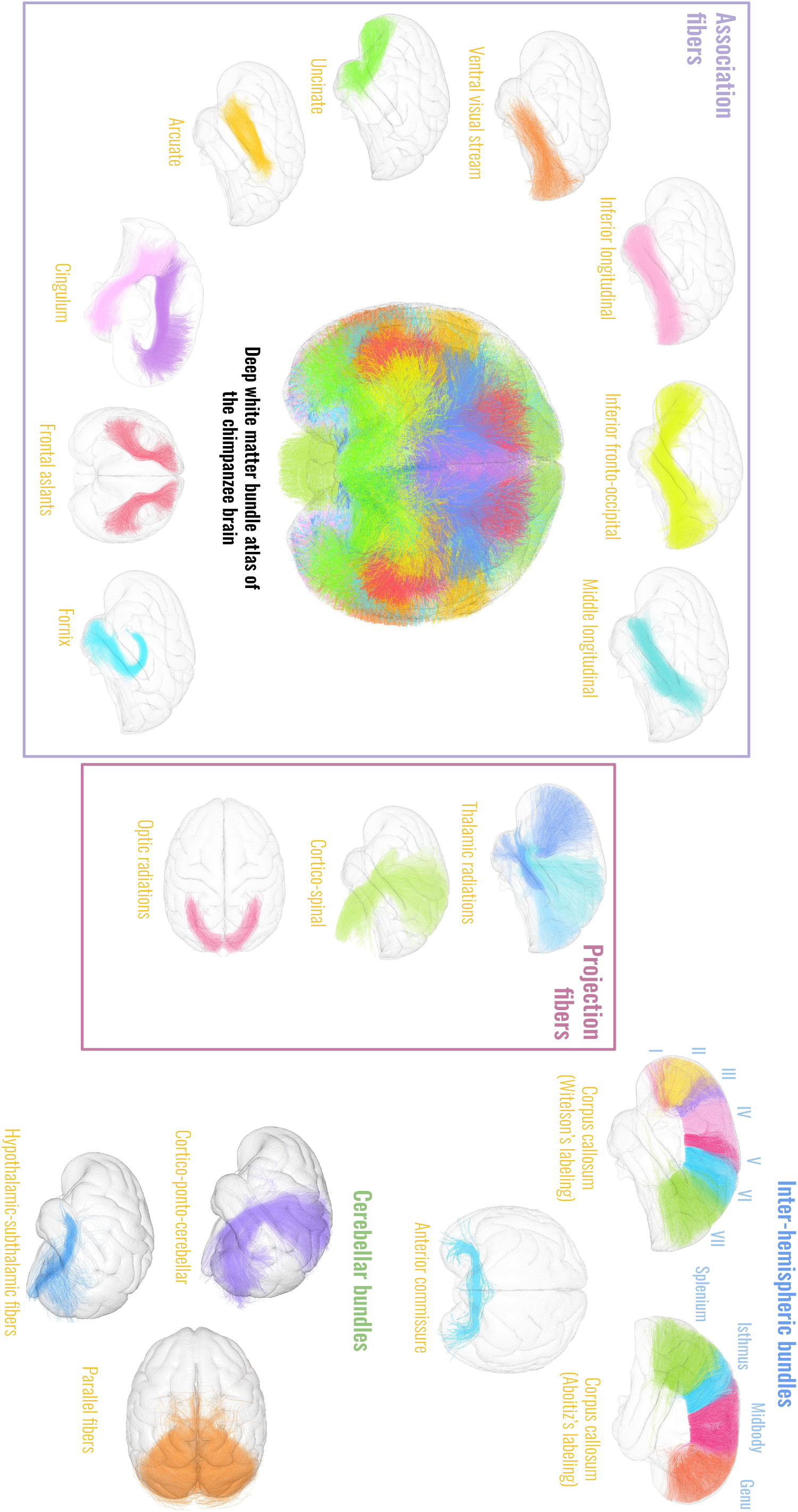
A new deep white matter fiber bundles atlas of the chimpanzee brain. 3D renderings of the 46 white matter bundles composing our long white matter atlas and including : the middle longitudinal bundle, the inferior longitudinal bundle, the arcuate, the frontal aslants, the uncinate, the fornix, the cingulum (dorsal and ventral), the motor and sensory ascending and descending fibers (cortico-spinal), the anterior commissure, the optic radiations, the ventral visual stream, the thalamic radiations (anterior, superior and posterior), the inferior fronto-occipital bundle; the cerebellar fiber components : the hypothalamic and subthalamic fibers, the parallel fibers, and the cortico-ponto-cerebellar fibers; the corpus callosum (CC) components : (left) the four subparts of the CC following Aboitiz labeling (splenium, isthmus, midbody, genu) and (right) the seven subparts of the Witelson labeling. (inf : inferior). White matter bundles are all represented with different colors and superimposed to the cortex pial surface computed from the DAVI130 atlas and rendered with transparency.

#### 3.1.1 Commissural fibers

The commissural fibers are composed of the anterior commissure (AC), the posterior commissure (PC) and the corpus callosum (CC). The AC crosses through the ventral part of the anterior wall of the third ventricle, rostrally to the columns of the fornix, and connects the two temporal lobes of the left and right hemispheres on their rostral and medial parts. The CC interconnects all the lobes of both hemispheres and is composed, rostrally to caudally, of the genu, midbody, isthmus and the splenium, as proposed by following the Aboitiz et al. 1992 convention and the 7 labels segmented parcellation approach advanced by Witelson 1989. Segmentation of these regions were performed in the mid-sagittal plane using the approaches described in Aboitiz et al. 1992 and Witelson 1989. As described in Goldstein et al. 2022 forceps minor and major are subparts of the CC and are present in this atlas under the different proposed nomenclatures. The forceps minor is composed of fibers passing thought the posterior part of the CC, connecting occipital lobes. They are considered here as the splenium, referring to Aboitiz segmentation, and to label 7 of the Witelson segmentation. The forceps major is composed of fibers passing through the anterior part of the CC, connecting frontal cortices. In this study they are considered as the genu, referring to Aboitiz segmentation, and to label 2 of the Witelson segmentation.

#### 3.1.2 Projection fibers

The projection fibers defined in our deep white matter bundle atlas are composed of efferent and afferent fibers. They correspond to pairs of ipsilateral white matter bundles including the: cortico-spinal tracts (CST), the thalamic radiations (anterior, superior, posterior) (TRa, TRs, TRp), the cortico-ponto-cerebellar tracts (CPCT), and the optic radiations (OR). The CSTs originate from the cortical motor areas, mainly from the precentral gyrus but also from the more rostrally located cortical areas (premotor cortex and supplementary motor area). Then, they pass through the corona radiata, the posterior limb of the internal capsule, and reach the anterior part of the brainstem (midbrain, pons and medulla oblongata). The TRs arise from the thalamus and reach different cortical areas: the frontal cortex for the TRas, the parietal cortex for the TRss, and the occipital cortex for the TRps. Projection fibers connecting the cerebral cortex and the pons or the cerebellar cortex were identified as the CPCTs. They originate from the superior frontal and parietal cortices and connect the cerebellum through the superior and middle cerebellar peduncles. It is important to note that a larger number of fibers connect the cerebellum through the ipsilateral rather than the contralateral peduncles. The ORs arise from the lateral geniculate nucleus, extend around the occipital horn of the lateral ventricle and project caudally to the primary visual cortex around the calcarine fissure in the occipital lobe.

#### 3.1.3 Association fibers

Among the association fibers reconstructed in our long white matter atlas are pairs of ipsilateral white matter bundles including: the inferior fronto-occipital fasciculi (IFOF), the inferior longitudinal fasciculi (ILF), the middle longitudinal fasciculi (MdLF), the ventral visual streams (VVS), the arcuate fasciculi (AF), the frontal aslant tracts (FAT), the uncinate fasciculi (Unc), cingulum (CG) (including ventral CGv and dorsal CGd) and the fornix (FX). The IFOFs are long fasciculi connecting the inferior frontal gyrus and the medial part of the occipital lobe, passing through the external capsule where their sections narrow. The ILFs connect the ventro-lateral part of the temporal poles and the inferior temporal gyri to the ventral part of the occipital lobes. These bundles run ventrally to the IFOFs, parallel to their paths in the occipito-temporal lobes. The MdLFs seems to originate in the superior temporal pole,run through the superior part of the temporal lobe and reaches the superior temporal gyri and the parietal lobe (the inferior parietal lobule/supramarginal gyrus). They are located dorsally to the ILFs and laterally to the IFOFs. The VVSs originate from the occipital lobe, run toward the temporal lobes ventrally to the MdLFs, laterally to the ILFs, and terminate in the middle temporal gyri. The AFs are curved white matter bundles that originate in the inferior frontal gyrus and extend caudally to the temporo-parietal junction. The FATs are shorter bundles located inside the frontal lobes, connecting the superior frontal gyri to the ventro-lateral part of the inferior frontal gyri.

The Uncs are curved white matter bundles which originate in the rostral part of the temporal lobes, bend upward rostrally from the anterior insula, and extend to the inferior frontal gyri and the orbito-frontal cortex. The CGs are divided into two subparts : the dorsal CGd and the ventral CGv parts. The CGv parts stem from the caudal parts of the cingulate gyri, passing through the parahippocampal gyri and reaching the hippocampus head. The CGd parts are located underneath the cingulate cortices, coursing dorsally along the CC to reach the medial prefrontal cortices. The FXs could also be partially reconstructed : they consist in C-shaped bundles that originate from the hippocampal head and extend caudally alongside the hippocampus. At the most caudal part of the hippocampal tail, the FXs bend medially and dorsally. Both FXs converge together at the medial surface of both hemispheres to form the posterior pillars of the fornices, which extend vertically, rostrally to the genu of the CC. They finally curve rostrally and run ventrally to the CC.

#### 3.1.4 Ponto-cerebellar fibers

Ponto-cerebellar fibers could also be identified including : the cortico-ponto-cerebellar tracts, described above, the parallel fibers and the hypothalamic/subthalamic fibers. The parallel fibers are numerous and closely packed fibers living in the cerebellar lobes, describing a parallel-like structure crossing the hemispheres fo the cerebellum.

### 3.2 Superficial white matter bundle atlas of the chimpanzee brain

One hundred and eleven pairs of cortical regions in the left hemisphere and 116 pairs of cortical regions in the right hemisphere present connections, corresponding to 422 white matter bundles (aggregating 76,845 fibers stemming from the individual tractograms) for the left hemisphere, and 400 white matter bundles (aggregating 74,091 fibers stemming from the individual tractograms) for the right hemisphere (see Figures 6, 8 and 9). Connections were found for all of the 38 cortical regions composing the Davi130 atlas hemispheres, which now allows the investigation of particular features of the chimpanzee cortical connectivity such as the central sulcus and the superior temporal sulcus (see figure 7).

**Figure 6:**
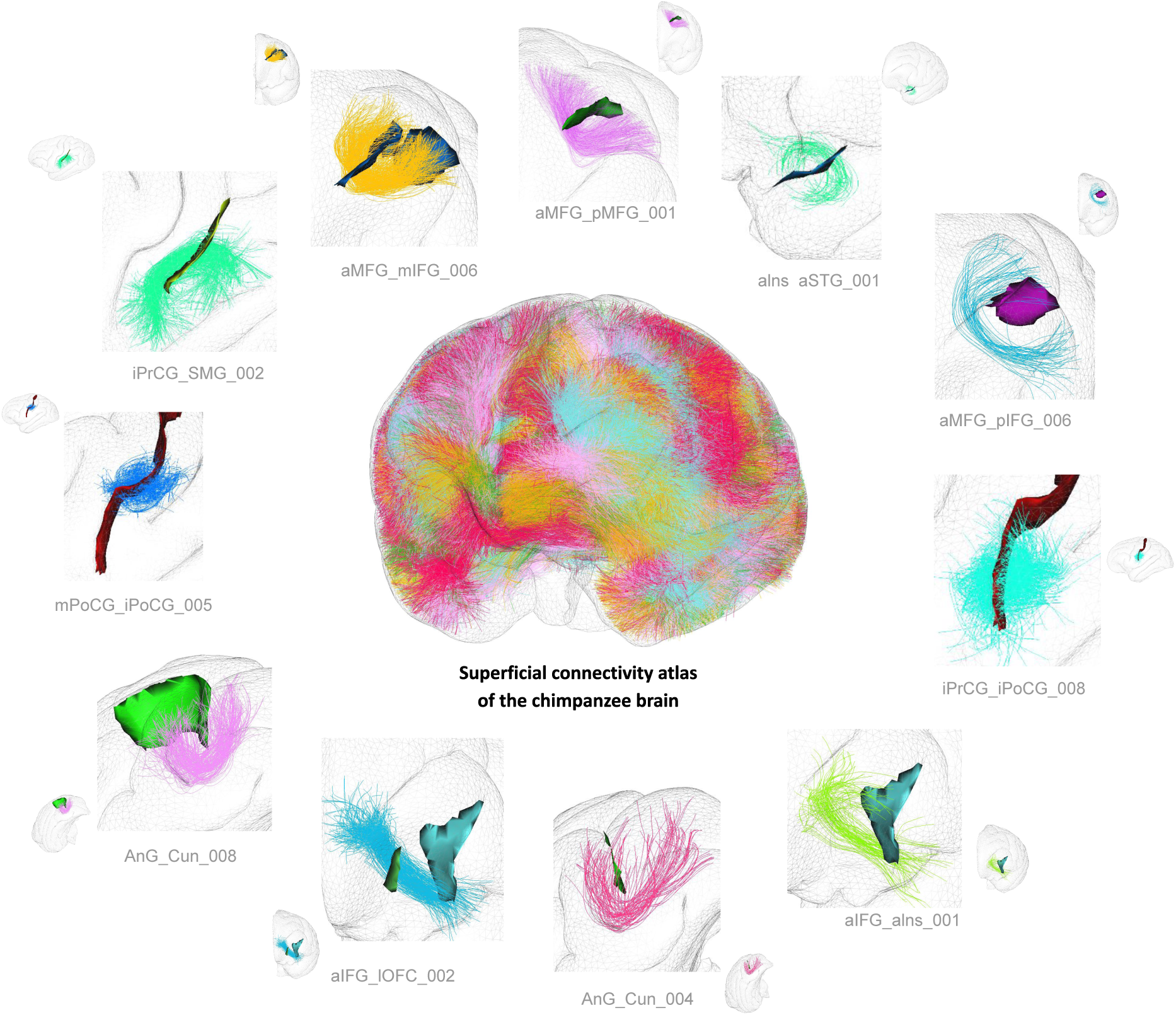
3D superimposition of the full set of white matter bundles composing the superficial white matter atlas and of the pial surface stemming from the Juna.Chimp template. (Center) Bundles composing the atlas, a different color is attributed to each of the 822 white matter bundles, surrounded by zooms over examples of superficial bundles composing the atlas, with the corresponding crossed sulci highlighting the coherence of the reconstructed fiber trajectories populating each bundle.

**Figure 7:**
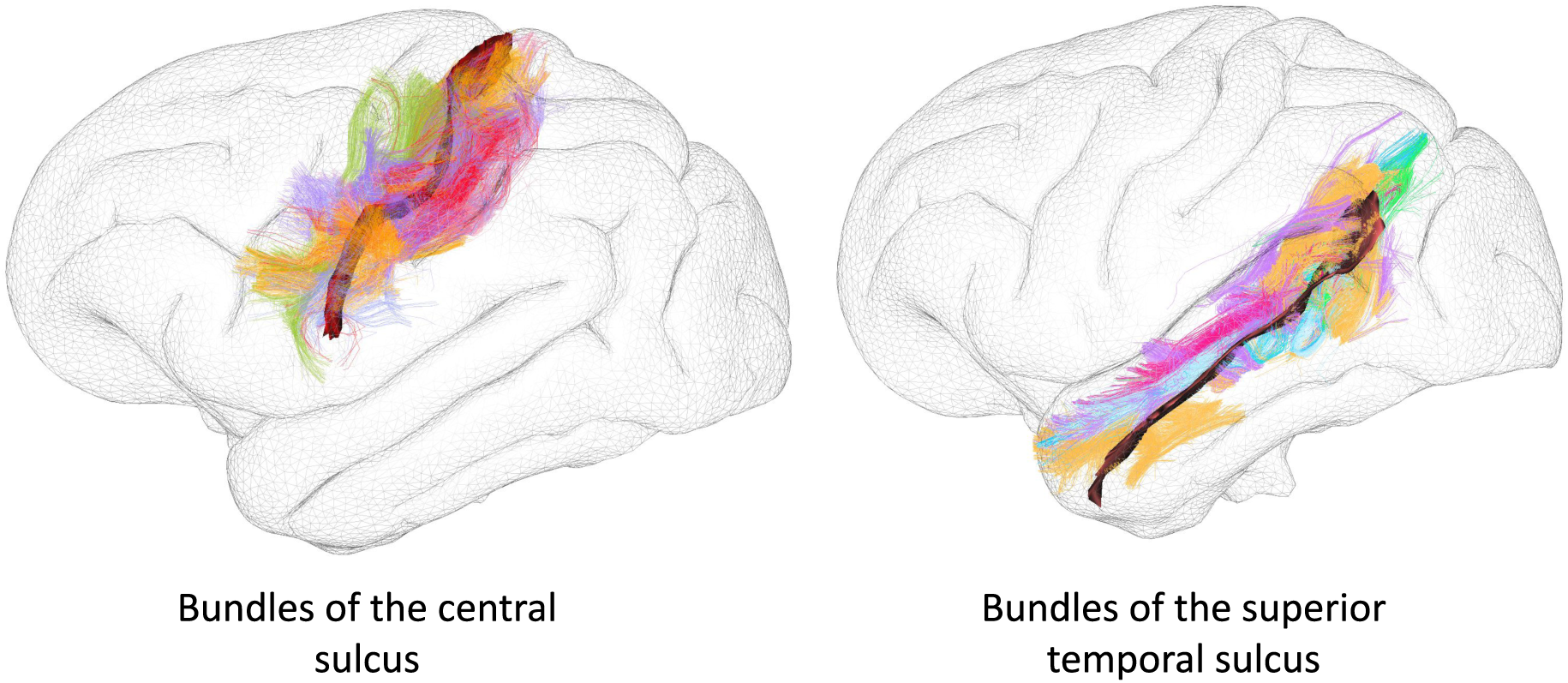
Short bundles surrounding the central sulcus (left) and the superior temporal sulcus (right) stemming from the superficial WM bundle atlas of the chimpanzee brain created in the frame of this work.

**Figure 8:**
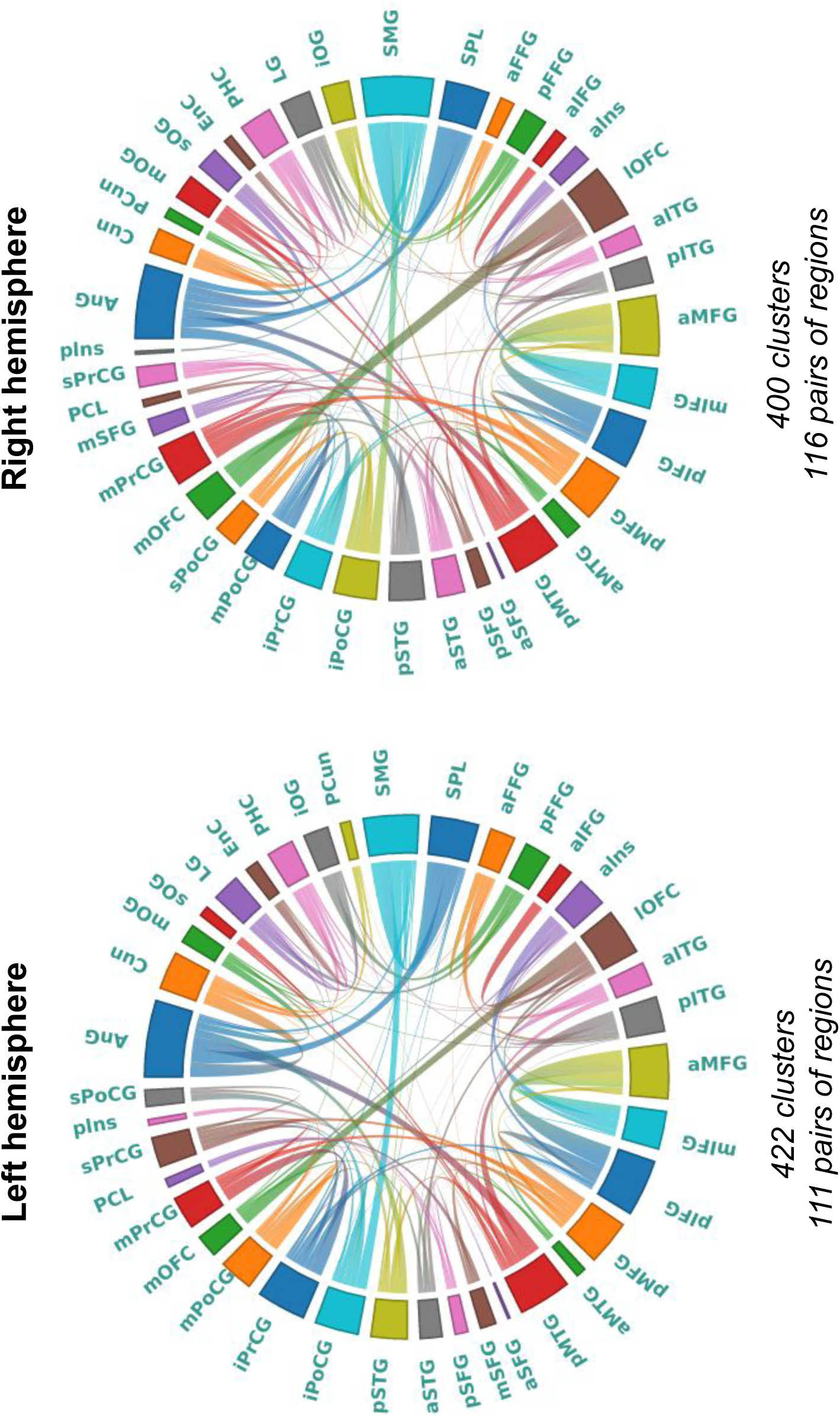
Graphical renderings of the 822 white matter bundles composing the superficial white matter connectivity atlas of the chimpanzee brain. (Left) 422 white matter bundles interconnecting 111 pairs of cortical regions from the left hemisphere as defined in the DAVI130 cortex atlas and (right) 400 white matter bundles interconnecting 116 pairs of regions from the right hemisphere

**Figure 9:**
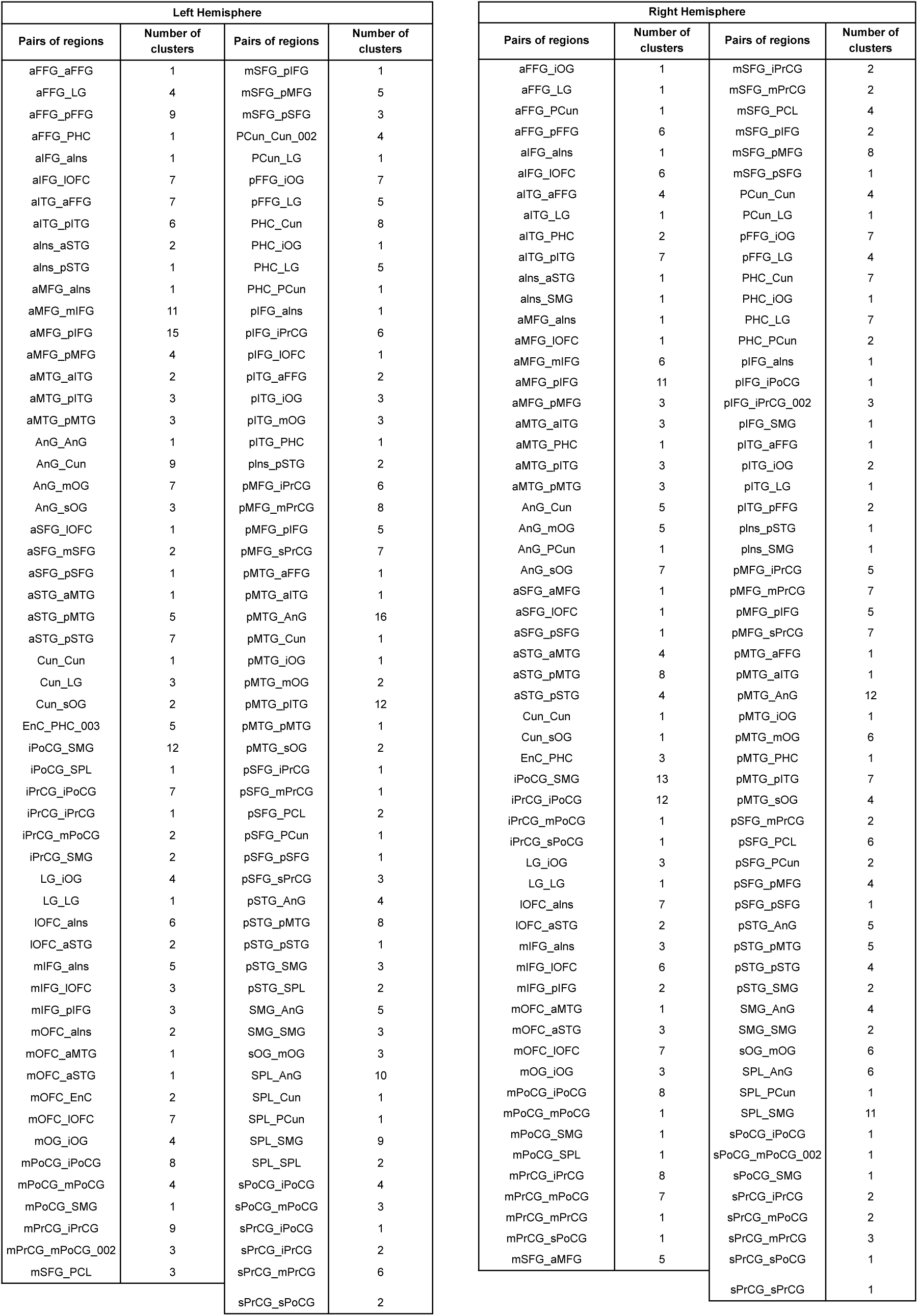
Table of the number of cluster per pair of regions

A significant variability of the number of fascicles (e.g. fiber clusters) originating from each cortical region was observed (see figure 8.). Notably, cortical areas AnG, pMTG and SMG depicted the largest number of fascicles in both hemispheres : AnG (LH : 56; RH : 45), pMTG (LH : 54; RH : 49), SMG (LH : 38; RH : 39). Some other cortical areas showed significant differences in connectivity between hemispheres, such as the mPoCG with 25 clusters in the left hemisphere versus 6 clusters in the right hemisphere. Cortical areas with a very low connectivity were also observed in both hemispheres such as the EnC, aSFG and plns areas : EnC (LH : 4; RH : 3); aSFG (LH : 4; RH : 3), plns (LH : 2; RH : 2). In the left hemisphere, the PCL regions notably contained few clusters (5), and in the right hemisphere, we noticed the sPoCG region to also display few clusters (6). The pair of regions sharing the highest number of clusters on both hemisphere were : pMTG_AnG (LH : 16; RH : 12), aMFG_pIFG (LH : 14; RH : 11), iPoCG_SMG (LH : 12; RH : 13). We also noticed that in the left hemisphere, the pair pMTG_pITG had 12 clusters.

After proceeding with the segmentation of the superficial white matter bundles of all the subjects of our cohort using our superficial atlas, we observed differences in the number of interconnections between regions (see figure 10). Indeed, the most interconnected regions were the angular (AnG) and supramarginal gyri (SMG). Some areas in the frontal lobe such as the middle frontal gyrus and superior parietal lobule also displayed a large number of connections. The motor and sensory motor areas, located along the central sulcus in each gyrus, presented a gradient of connectivity with a higher number of connections present in the inferior, pre- and postcentral gyri compared to the middle and superior pre/post central gyri. A higher number of superficial white matter bundles was observed in the motor regions (e.g. precentral gyrus) compared to the somatosensory regions (e.g. postcentral gyrus). The temporal lobe also depicted a higher density of connections in the posterior temporal gyrus (including the posterior superior, posterior middle and posterior inferior gyri) compared to the anterior temporal gyrus (containing the anterior superior, anterior middle and anterior inferior gyrus). In both hemispheres, the posterior middle temporal gyrus was the most interconnected gyrus in the temporal lobe.

**Figure 10:**
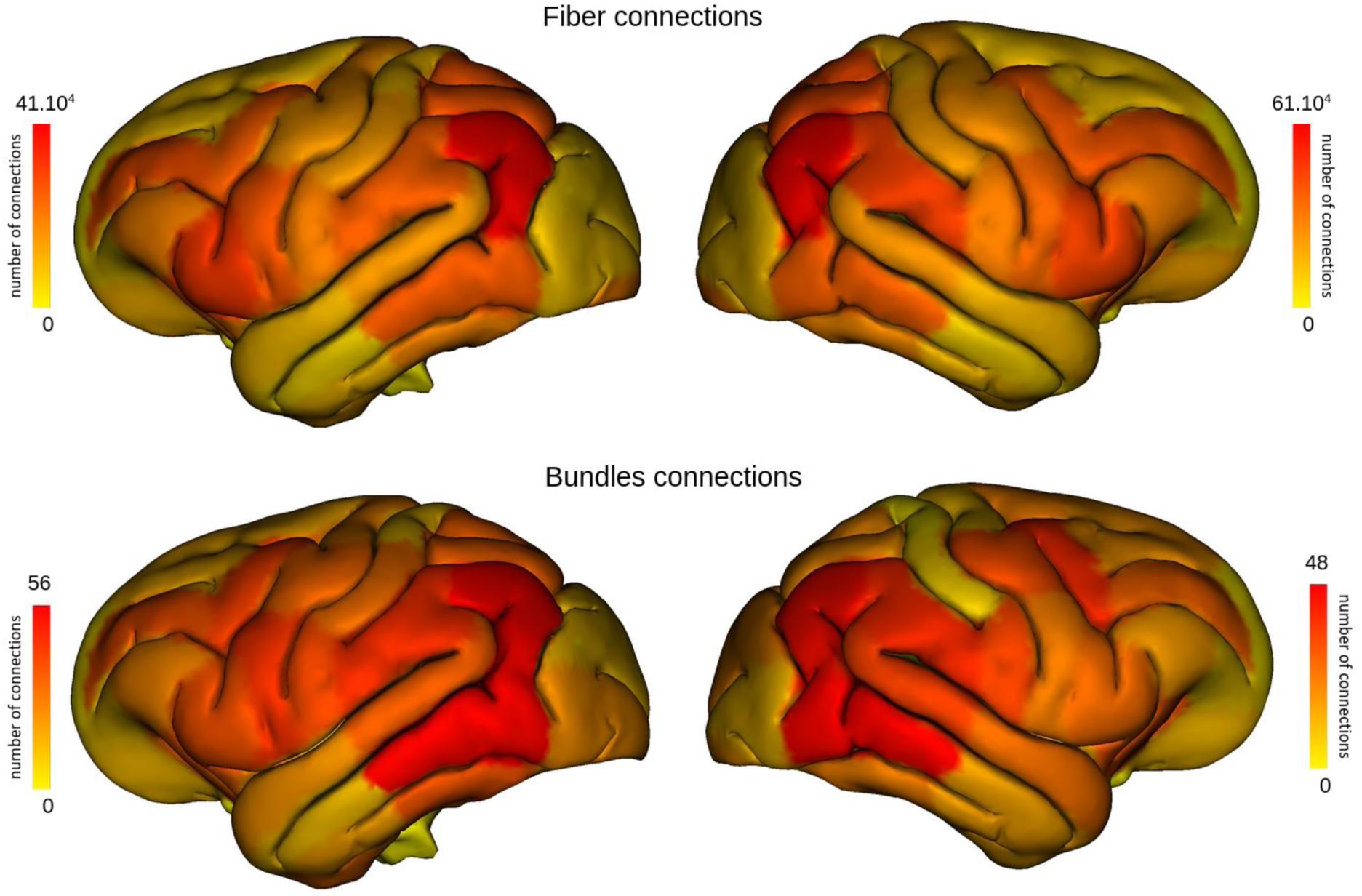
Average density maps of superficial streamlines (top) and white matter fascicle clusters (bottom) for the left and the right hemispheres resulting from the automatic segmentation of the superficial white matter bundles of all chimpanzees using the established superficial white matter atlas.

Within each hemisphere, we found a diversity in the shapes of the short fibers (see Figure 11). Indeed, symmetrically between the left and the right hemisphere, we observed : “U’”-shaped fibers, “V”-shaped fibers and “L”-shaped fibers. The “U” fibers were located in all lobes of the brain, the “V” fibers were mostly found in the superior frontal areas and superior temporal lobe, and the “L” fibers depicting a flatter shape could be seen running ventrally of each hemisphere.

**Figure 11:**
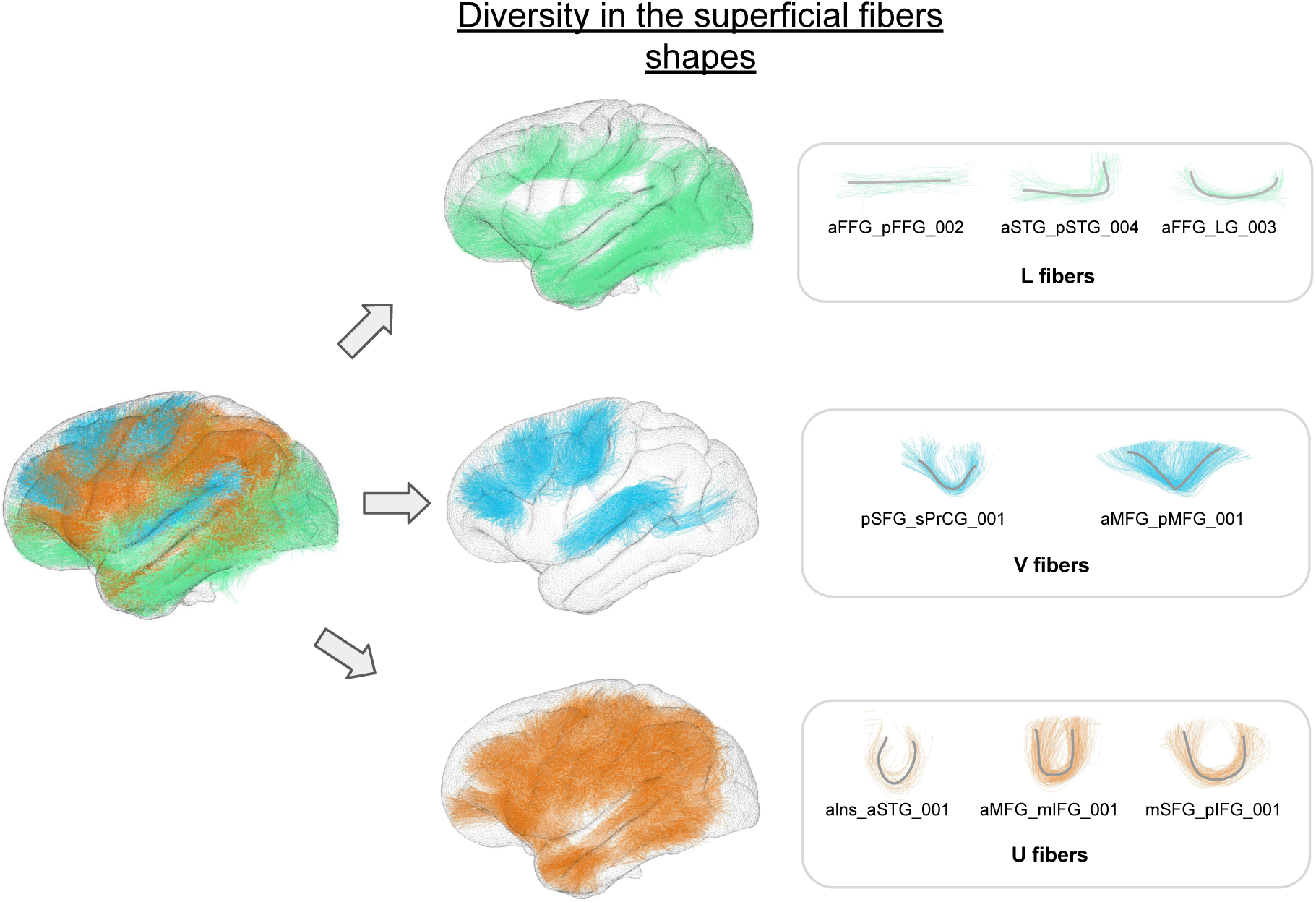
Diversity in the superficial fibers shapes. Representation of the distribution of the superficial white matter bundles with respect to 3 identified superficial bundle shapes: “L” bundle shape, “V” bundle shape and “U” bundle shape.

## 4 Discussion

In recent years, there has been renewed interest in mapping the anatomical and functional organization of the chimpanzee brain, whose proximity to human species may provide the key to understanding its commonalities and differences with the human brain. Structural connectivity is an essential part of the information for understanding the organization of functional networks and some studies already pointed out exciting outcomes concerning the chimpanzee brain neuro-anatomy(K. Bryant et al. 2020, K. L. Bryant et al. 2021, Hecht et al. 2015, K. A. Phillips and Hopkins 2012, Rilling et al. 2011, Roumazeilles et al. 2020,Mars et al. 2019) and mostly focusing on deep white matter bundles. To our knowledge, none of the previous dMRI studies in chimpanzees has addressed the organization of the superficial connectivity of the chimpanzee brain on a global scale although it is of major interest for a better comprehension of high level cognitive functions, largely unknown in this species. Thus, our findings, particularly on the superficial bundles, represent a unique contribution to the field. In this study, we have designed a novel deep white matter atlas, composed of 46 long/deep white matter bundles and have proposed a complete subdivision of the corpus callosum. This atlas was extended with a new superficial white matter atlas composed of 822 sub-cortical white matter bundles which establishes, to our knowledge, the first superficial white matter bundle atlas of the chimpanzee brain. The methodology used to build this atlas was different in essence from the one proposed in (K. Bryant et al. 2020) using a two-step method that does not require predefined regions of interest at any time, thus making it more general and less operator dependent.

### 4.1 Deep white matter bundle atlas of the chimpanzee brain

The deep structural connectivity atlas presented in this work consists of 46 bundles whose segmentation relies on a significantly different segmentation method than the approach used previously (K. Bryant et al. 2020) to establish a deep white matter bundle (DWMB) atlas. In particular, the approach of this work did not require the definition of specific seed ROIs and target ROI, which typically involves a preliminary segmentation process or anatomical assumptions. The clustering approach proposed in this study is fully automatic, observer independent, and only relies on the final labeling of segmented fascicles at the very end of the process. All the well-known commissural, projection, association and cerebellar white matter bundles could be easily segmented to compose the white matter bundle atlas. Some previous observations made on chimpanzee fiber bundles were consistent with the results we found in our atlas. For instance, the frontal aslant tract (FAT), which is also known for its role in language and speech (Dick et al. 2019) was clearly identified and strikingly resembles its homologous tract in humans. In the human species, the left FAT seems to support speech and language functions, whereas the right FAT role seems to be the support of executive functions, inhibitory control and conflict monitoring (La Corte et al. 2021). In this study, no population-level lateralization in fiber density was demonstrated, however, the FAT had a non-negligible thickness and was present in all subjects. The fact that this fasciculus resembles its human homologue is also important considering the major differences in the frontal lobe organization of the chimpanzee compared to humans (Alatorre Warren et al. 2019, Amiez et al. 2019). The presence of the arcuate fasciculi in non-human primates has been a subject of debate in the scientific community (Becker et al. 2022, Balezeau et al. 2020). Our approach was able to reconstruct the arcuate fasciculi (AF) with a trajectory that appears as similar with previous studies (K. Bryant et al. 2020) however with some nuances. While the AF projections are connecting the inferior frontal gyrus to the inferior parietal lobe and the superior temporal gyrus, they appear to be thin and less extended than what could be expected compared with humans. We observed present but weak connections in the superior part of the temporal lobe. The results presented here also confirm the study from (James Rilling et al. 2008) showing an arcuate fasciculus with less temporal extensions in the chimpanzee brain compared to the human brain. This particular temporo-parietal area reached by the AF corresponds to the planum temporale and is identified as having a key role in language and speech in the human brain (Shapleske et al. 1999). We did not specifically recover the superior longitudinal fasciculus III (SLF-III), whereas other studies revealed its presence in the chimpanzee brain (K. Bryant et al. 2020, Hecht et al. 2015). However, due to the proximity and possible overlap with the AF, the fiber clustering may have merged the two bundles into one. A finer tuning of the distance threshold will be tested in the future to see if we can recover the SLF and AF separately. Further investigating the language pathways, we clearly identified the IFOF, which has also been the subject of interrogation regarding its presence and role in the non-human primate brain (K. Bryant et al. 2020, Roumazeilles et al. 2020). In our cohort, it displayed relatively strong connections between superior and inferior endings in the prefrontal cortex and the occipital lobe, and is a thick and consistent bundle. The middle longitudinal fasciculus (MdLF) could be reconstructed laterally to the IFOF. This bundle is controversial since its morphology and function are still poorly understood nowadays in humans (Latini et al. 2021). It has been described as connecting the temporal, parietal and occipital lobes. They are located dorsally to the ILFs. In the case of the study presented here, the MdLF originates in the superior temporal pole, runs through the superior temporal lobe and reaches posterior superior regions of the temporal lobe as well as parietal regions in the inferior parietal lobule (IPL)/supramarginal gyrus (SMG), as described in (Seltzer and Pandya 1984) in the macaque brain. The MdLF endings seem to reach the ILF or SMG without going further in the superior parietal lobule or the occipital lobe, consistent with what was described in (K. Bryant et al. 2020) in chimpanzees. However, we have noticed that a portion of the posterior extremity of the MdLF presents a decussing showing connections with some posterior middle temporal regions. This particular aspect of the bundle has not been previously described. The other white matter fiber bundles do not depict any particular morphological difference except the effect due to the difference between chimpanzee and human postures (mostly quadrupedal versus bipedal) as well as brain rearrangements oc-curring during the hominin evolution (Alatorre Warren et al. 2019). This can be observed for the CST and CPCT. Indeed, bundles connecting cortical areas with the basal ganglia or with the brainstem displayed an adaptation due to the posture anatomy. The corpus callosum bundle, subdivided into its genu, isthmus, midbody and splenium components, confirms the study from Phillips and Hopkins (K. A. Phillips and Hopkins 2012) as its organization resembles the one of humans. The cerebellar components reconstructed in this study for the chimpanzee brain are structurally similar to the human ones. Indeed, concerning the cortico-ponto-cerebellar tracts, as exposed in (Palesi et al. 2017), fibers originate not only in motor areas but also in more frontal and parietal areas, and connect cerebellar hemispheres. It is nevertheless important to note that this bundle is usually described to be composed of fibers reaching contralateral cerebellar hemispheres. In the frame of this study, we only saw connections between cortical regions and ipsilateral cerebellar hemispheres, which have been shown to also exist in humans (Karavasilis et al. 2019). We can propose different hypothesis to explain this phenomenon: first, this highlights limitations in the methodology used to build the atlas, especially concerning the intra-subject fiber clustering. Indeed, the second step consists in splitting the groups of fibers with respect to the regions they belong to (left hemisphere, the right hemisphere the interhemispheric and the brainstem/cerebellar regions). This could cause an artificial break in tracts that would be considered mostly belonging to the right and left hemispheres but that would also reach the cerebellum, causing a mismatch of the fiber clusters corresponding to this bundles in ROI attribution of the clusters building, not considering fibers reaching the contralateral cerebellum hemisphere but only taking into account the fibers comprised on the same side. An easy solution to this issue consists of first splitting the cerebellum into its two hemispheres, then creating two further regions, one comprising the right hemisphere and left cerebellum, and the other one comprising the left hemisphere and right cerebellum. Applying our clustering approach to these two new composite regions should better highlight clusters corresponding to the cortico-ponto-cerebellar target white matter bundles. Another possible hypothesis, completing the previous one, is that the number of fibers connecting one hemisphere and reaching the contralateral cerebellar hemisphere are much less numerous in the chimpanzee brain compared to what have been previously described in the human brain. However this hypothesis seems unlikely. Since the study presented here is the first one depicting CPCT in chimpanzees, this would need to be further investigated, whether using ROI or relaunching a fiber clustering considering the whole brain.

### 4.2 Superficial white matter bundle atlas of the chimpanzee brain

Superficial white matter bundles (SWMB) are mostly known to be short sub-cortical association fibers or U-fibers, connecting surrounding gyri (Meynert 1885). In comparison to deep white matter bundles extensively described in the literature, SWMB were poorly described in the literature due to their small lengths and sections and to their sub-cortical positioning making their dissection difficult using Klingler’s method (Klingler and Ludwig 1956). Fiber tracking tools based on diffusion MRI data have been used to explore this superficial connectivity for little more than a decade, albeit with intrinsic limitations related to the low resolution of MRI data, the ill-posed nature of the problem of local inference of fiber directions and fiber trajectories by tractography, but also because of the large number of superficial bundles populating the subcortical region. However, diffusion-based fiber tractography and clustering methods have been successful to reconstruct some of the human superficial white matter bundles (M. Guevara et al. 2020, Zhang et al. 2014, Labra Avila 2020). SWMB results from brain development and continuously evolves during the maturation process (O. R. Phillips, Clark, et al. 2013, M. Wu et al. 2014). Further, they are known to be affected during brain aging (Nazeri, Chakravarty, Felsky, et al. 2013, Nazeri, Chakravarty, Rajji, et al. 2015) and in brain disorders (O. R. Phillips, Joshi, et al. 2016, O. Phillips et al. 2016). Few studies have investigated the superficial connectivity of non-human primates (Oishi et al. 2011). The first superficial white matter bundle atlas of the chimpanzee brain established in the frame of this study is a further demonstration of their existence in the chimpanzee brain and might contribute to the identification of commonalities and differences of the superficial connectivity between chimpanzees and humans, in order to better understand the singularity of the human brain.

While K. Bryant et al. 2020 also addressed superficial connectivity describing a few short white matter bundles, the present study offers a more exhaustive exploration of the superficial structural connectivity, aiming at proposing a complete mapping of the short white matter bundles, despite remaining mostly exploratory and descriptive. Among the hundreds of superficial white matter bundles reconstructed in this study, we did observe a slightly higher number of bundles populating the left hemisphere compared to the right hemisphere. However, the right hemisphere seems to contain more interconnected regions. When looking at homologous superficial white matter bundles between hemispheres (eg linking two homologous regions in each of the two hemispheres), their number of fibers are of the same order of magnitude thus sharing a similar connectivity profile. The regions depicting the highest number of connections are situated in the parietal lobe and on the ventral portion of the frontal lobe. As expected, most SWMB depicted a ‘U’ shape, but we did observe some other shapes such as ‘V’, ‘L’ or flattered shapes. While the ‘U’-shaped fibers seem to be situated homogeneously centrally in the lateral part of the brain, the ‘V’-shaped fibers were mostly found on the dorsal part of the brain. The ‘L’ fibers seem to be part of the longest superficial white matter bundles and are mainly located on the ventral part of each hemisphere. Because we separated SWMB from DWMB using a simple threshold over their fiber length, the SWMB atlas also includes fiber clusters belonging, for instance, to the optic radiations or to subparts of the frontal aslant tracts. As no consideration strictly compels their classification into long or short bundles, we considered them in both atlases. It is also the case concerning the vertical occipital fasciculi, identified in the SWMB as “Left_sOG_mOG” and “Right_sOG_mOG”, subdivided into 3 clusters on the left hemisphere and 6 clusters in the right hemisphere.

### 4.3 Limitations

As with any imaging study, the current work is subject to biases due, on the one hand, to the limitations of the acquisition methods and, on the other, to the limitations of the analysis methods employed. Diffusion MRI is known to be particularly sensitive to acquisition noise, given the limited signal-to-noise ratio associated with the very principle of attenuation of the MRI signal by the diffusion process of water within brain tissues. Moreover, tractography techniques are known to induce numerous artifactual fiber trajectories, which need to be ruled out using correction methods that take anatomical prior knowledge into consideration. While it is easy to define such a priori for deep connectivity, which is relatively well established, it seems more difficult to define them for superficial connectivity, which remains largely unexplored today, particularly in chimpanzees. An additional difficulty in studying the anatomical connectivity of the chimpanzee brain is the relative difficulty of access to ex vivo anatomical brain specimens, which greatly limits the possibility of validating results obtained in vivo from MRI, for example using Klinger’s dissections or microscopic approaches such as Polarized light imaging (PLI) or Optical coherence tomography (OCT).

The validity of the tracts presented here rely considerably on the fiber tracking method used. In the frame of this study, the regularized deterministic streamline approach of (Perrin et al. 2005) was employed which allows to efficiently deal with fibre crossing configurations, and therefore significantly reduce the number of false positives as reported in (Sarwar, Ramamohanarao, and Zalesky 2019), thus leading to a better reconstruction of the whole brain tractograms from all the subjects, from which the clusters derive. However, studies have also shown that in the case of higher fiber complexity, deterministic tractography may not be sufficient and be very sensitive to the principal estimated direction and noise (Descoteaux, Deriche, et al. 2008, Jones 2010). We could alternatively use probabilistic or even better global approaches, but we have decided at a first glance to consider the regularized deterministic approach. We intend investigate probabilistic and global approaches in the future to study their impact on the reconstructed white matter bundle atlases. The millimeter resolution of the anatomical and diffusion MRI datasets we used is obviously a limiting factor, but the ENPRC biobank remains today the largest cohort of imaging data acquired on the chimpanzee brain in vivo. The emergence of new high-end connectome gradients and high field MRI scanners might change the future of the exploration of the chimpanzee brain. It should offer the possibility to scan chimpanzees at a much higher spatial resolution (from submillimer scale to mesoscale) and to significantly improve the SNR of the dMRI dataset, which should allow to more accurately map the endings of the connections entering the cortical ribbon and therefore more reliably reconstruct superficial white matter bundles.

However, in its current implementation, this two-step fiber clustering algorithm proposes a robust approach to the inference of white matter bundles, automatically and pretty efficiently discarding outliers. This was indeed the case with only a few remaining artifact fascicles, mostly located superficially at the boundary of the brain mask in regions where the mask was not perfectly delineated. The effect of using a two-step fiber clustering together with a fiber registration to a template may have introduced some inter-individual fiber shape differences that may explain why some of the bundles could not be clearly identified, such as the SLF I and II (and supposedly the distinction between the arcuate and the SLF III) as well as the auditory radiations, contrarily to studies from Hecht et al. 2015 and K. Bryant et al. 2020. Furthermore, due to the separation of the two hemispheres during the first intra-subject fiber clustering, some fibers connecting one hemisphere to the other might have been missed during the reconstruction process as explain in the discussion section concerning the cortico-ponto-cerebellar tracts. It remains challenging to make the distinction between longest (deep) and short (superficial) white matter bundles since this dual definition is on the order of ontological considerations without a real split between the two concepts. It is fair to say that in this study we’ve taken it upon ourselves to say that the superficial bundles are cortico-cortical connections that are not part of the association bundles presented in the deep white matter atlas. We based ourselves on the threshold given by the cumulative histogram of fiber lengths stemming from all the subjects tractograms, resulting in a drop of the number of fibers at 50 millimeters. We then performed multiple tests in pairing the white matter bundles using different thresholds, starting from the consideration of all the fibers and pairing them to the different labeled contained in the DAVI130 atlas. We reduced the threshold as long as we would exclude the major deep white matter bundles and found the threshold to be between 50 and 40 millimeters. We kept 50mm to try to ensure that most of the superficial fibers would be taken into account but this still rely on a empirical choice, and thus this as to be taken into consideration with respect to the atlas presented here. In order to properly validate the data presented here, the same comparable investigation needs to be performed on human data.

### 4.4 Perspectives

The current work has allowed us to establish a complete mapping of the deep and superficial connectivity in chimpanzees. If the validation of the large white matter bundles is relatively straightforward, the validation of the superficial bundles described in the atlas resulting from this study is more subtle, due to the lack of a reference in the matter. Therefore, a validation phase is necessary, which will require the acquisition of a few anatomical brain specimen scanned *ex vivo* using meso or micro scale imaging methods such as ultra-high field diffusion MRI or polarized light imaging high resolution images will allow for a better assessment the bundles described in our superficial white matter bundles atlas. Having the possibility to perform functional MRI (fMRI) studies with advanced stimuli would also contribute to better identifying the functional networks to which they belong too and to better understand their functional role. The observation of a limited number of shapes characterizing the geometry of short bundles compels us to further study the link between the shape of a short bundle and the sulco-gyral geometry of the cortical areas it connects, but also to look at the distribution of these connection shapes on the surface of the cortex. Lastly, the existence of atlases of superficial connectivity in both humans and chimpanzees opens the way to comparative studies to identify their key differences and to correlate these differences in connectivity with behavioral differences within and between the two species.

## 5 Acknowledgments

We thank the Ile de France Region and the Blaise Pascal International Chair of Excellence awarded to Pr. William Hopkins for funding this work. Additional support was provided by NIH grant NS-092988 (support for the National Chimpanzee Brain Resource).

## 6 Data availability

The superficial white matter bundles atlas and the deep white matter bundle atlas are fully available at :

- https://zenodo.org/record/7147503 - Ginkgo Chauvel’s left and right superficial white matter atlas of the chimpanzee brain;
- https://zenodo.org/record/7147789 - Ginkgo Chauvel’s deep white matter atlas of the chimpanzee brain.

## 8 Code availability

The entire pipeline used to design the structural connectivity atlases is freely available on Framagit (https://framagit.org/cpoupon/gkg).

