## Supplementary Figures and Tables for "*In vivo* mapping of the deep and superficial white matter connectivity in the chimpanzee brain"

### 1 Supplementary information

#### 1.1 SI 1 - Formula of the corrected distance chosen to compute fascicle clusters

The objective of this formula is to correct the evaluation of the symmetric pairwise distance between 2 centroids representing 2 individual fascicles in order to relax the constraints over the centroid distance when addressing long fascicles while keeping the distance unchanged for superficial (short) fascicles.

Let $C_1$ and $C_2$ be two centroids stemming from the intra-subject fiber clustering step. The centroids are represented by $N_p$ control points $\{P_{c_1}(i)\}$ and $\{P_{c_2}(i)\}$ (21 control points in our case). Let $d_{pairwise}(c_1, c_2)$ be the symmetric pairwise distance between centroids $c_1$ and $c_2$.

It yields:

$$d_{pairwise}(c_1, c_2) = min(\sqrt{\sum_{i=0}^{N_p-1}(P_{c_1}(i) - P_{c_2}(i))^2}, \sqrt{\sum_{i=0}^{N_p-1}(P_{c_1}(i) - P_{c_2}(N_p - i))^2}) \quad (1)$$

A corrected distance was implemented to relax constraints over the distance function when the minimum centroid length $L_{min} = min(length(C_1); length(C_2))$ increases in order to take into account the behavior of centroids showing more fanning configurations at their extremities in long fascicles in comparison to superficial ones:

$$d_{corrected}(c_1, c_2) = d_{pairwise}(c_1, c_2) - N_f(t_1T_1 + 1.3t_2T_2 + 1.6t_3T_3) \quad (2)$$

where:

$t_1 = 1$ if $L_{min} \geq l_{min}$ or $0$ otherwise

$t_2 = 1$ if $L_{min} \geq l_{min} + (l_{max} - l_{min})/3$ or $0$ otherwise

$t3 = 1$ if $L_{min} l_{min} + 2(l_{max} - lmin)/3$ or $0$ otherwise

$T_1 = \frac{min(L_{min}, l_{min} + (l_{max} - l_{min})/3) - l_{min}}{l_{max} - l_{min}}$

$T_2 = \frac{min(L_{min}, l_{min} + 2(l_{max} - l_{min})/3) - (l_{min} + (l_{max} - l_{min})/3)}{l_{max} - l_{min}}$

$T_3 = \frac{min(L_{min}, l_{max}) - (l_{min} + 2(l - max - lmin)/3)}{l_{max} - l_{min}}$

The normalizing factor $N_f$ controls the degree of relaxation of the distance, being typically chosen to 26 mm in our case for chimpanzees and 6 mm for humans. $l_{min}$ and $l_{max}$ define the lower and upper fiber lengths that were defined from the input subject tractograms (chosen to be 7mm and 133mm respectively) in order to remove the smallest and longest artifactual fibers.

#### 1.2 SI 2 - Optimization of the normalization factor of the corrected distance $N_f$ and of the affinity maximum distance $d_{max}$

In order to maximize the number of generated inter-subject fascicle clusters, the distance normalization factor $N_f$ and the maximal centroid distance threshold $d_{max}$ are optimized using a grid search algorithm for 50 values of $N_f$ uniformly sampled within the [2;100] range and for 17 values of $d_{max}$ uniformly sampled within the [2mm;34mm] range. This pre-evaluation is performed on a 10-subject subset of the population, which ensures to sufficiently take into account the cross-subject variability and allows to significantly speed-up the computation time. As shown in figure 1, the representation of the number of fascicle clusters with respect to $N_f$ and $d_{max}$ depicts a convex landscape which facilitates the identification of the optimal setting $(N_f, d_{max})_{opt} = (26, 18mm)$.

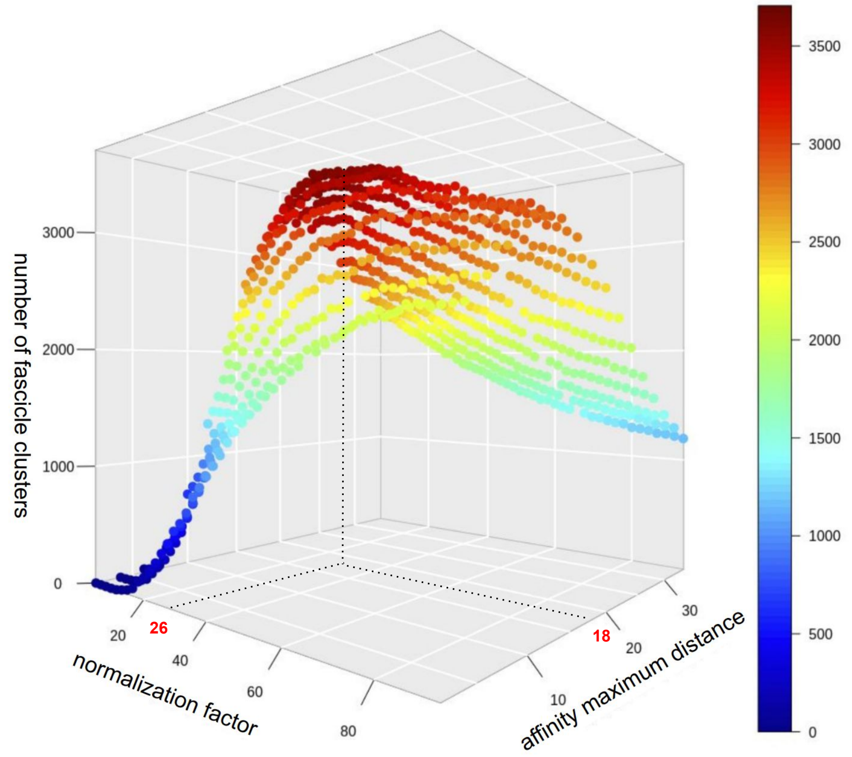


*Figure 1: Optimization of the normalization factor of the corrected distance $N_f$ and of the affinity maximum distance $dmax$ for identification of the optimal setting : $(N_f, d_{max})_{opt} = (26, 18mm)$. The maxima was reached with 3705 fascicle clusters.*

#### 1.3 SI 3 - Long white matter bundles composing the atlas

The **commissural fibers** are composed of the anterior commissure (AC) and the corpus callosum (CC) (see figure 2 and 3).

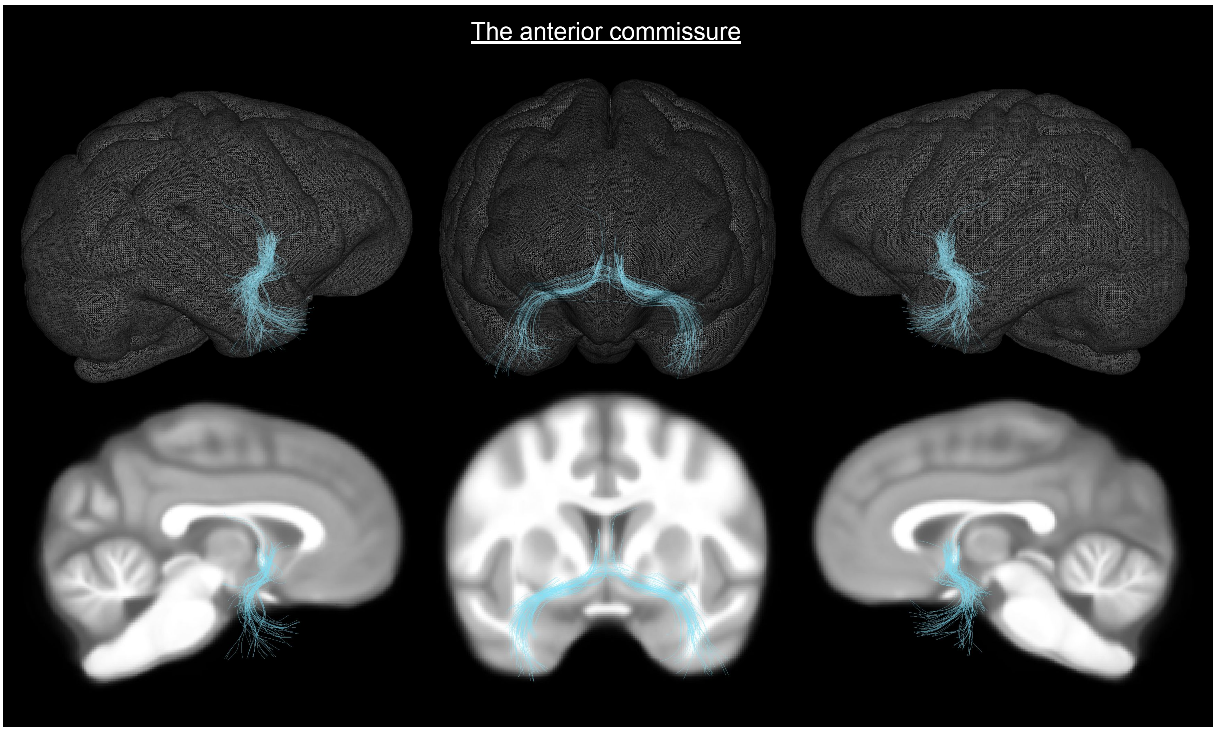


*Figure 2: A commissural fiber : the anterior commissure. (Top) tracts superimposed on a 3D mesh of the Juna.Chimp template; (Bottom) tracts superimposed on the T1-weighted anatomical image of the Juna.Chimp template. left sagittal, coronal, and right sagittal views.*

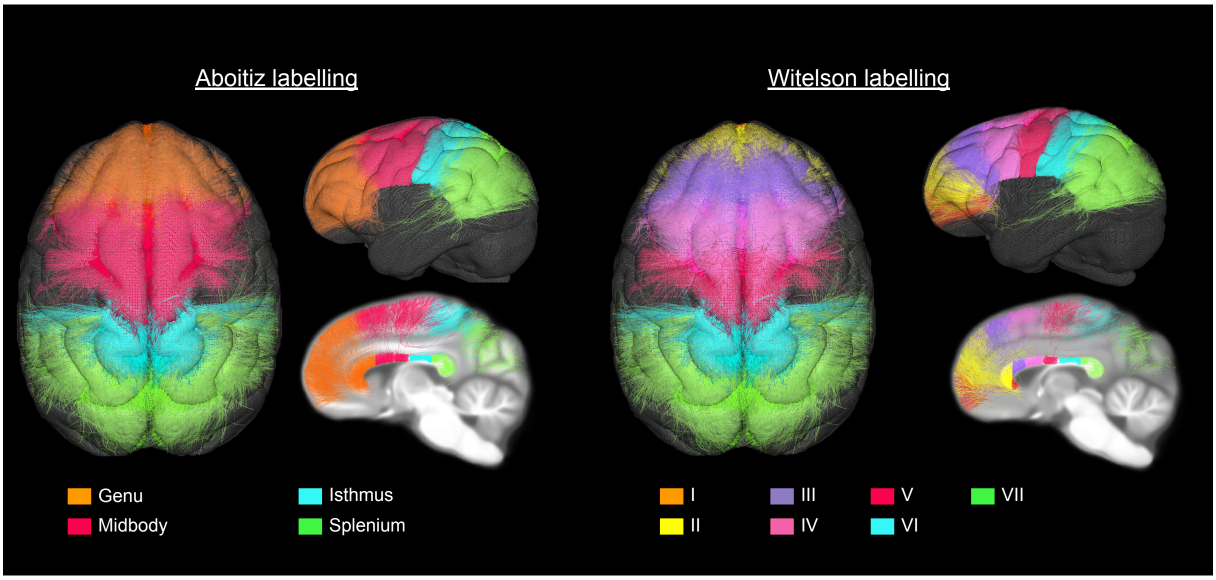


*Figure 3: A commissural fiber : the Corpus Callosum. Left : Aboitiz labelling of the CC displayed on the Juna.Chimp template mesh and anatomical image ; right : Witelson labelling of the CC displayed on the Juna.Chimp template mesh and anatomical image.*

**The projection fibers** proposed in our long white matter bundle atlas are composed of efferent and afferent fibers. They correspond to pairs of contralateral white matter bundles including : the cortico-spinal tracts (CST), see figure 4, the thalamic radiations (anterior, superior, posterior), see figure 5, the cortico-ponto-cerebellar tracts (CPCT), see figure 6, and the optic radiations (OR), see figure 7.

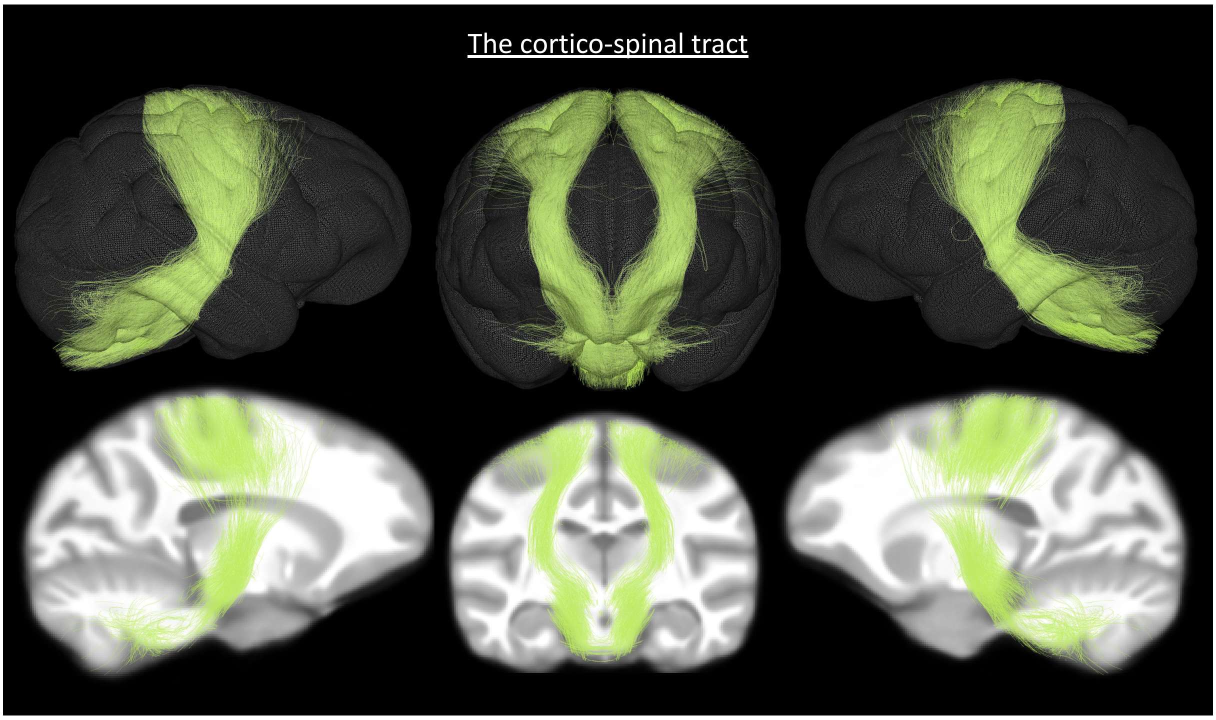


*Figure 4: The cortico-spinal tracts. (Top) tracts superimposed on a 3D mesh of the Juna.Chimp template; (Bottom) tracts superimposed on the T1-weighted anatomical image of the Juna.Chimp template. left sagittal, coronal, and right sagittal views.*

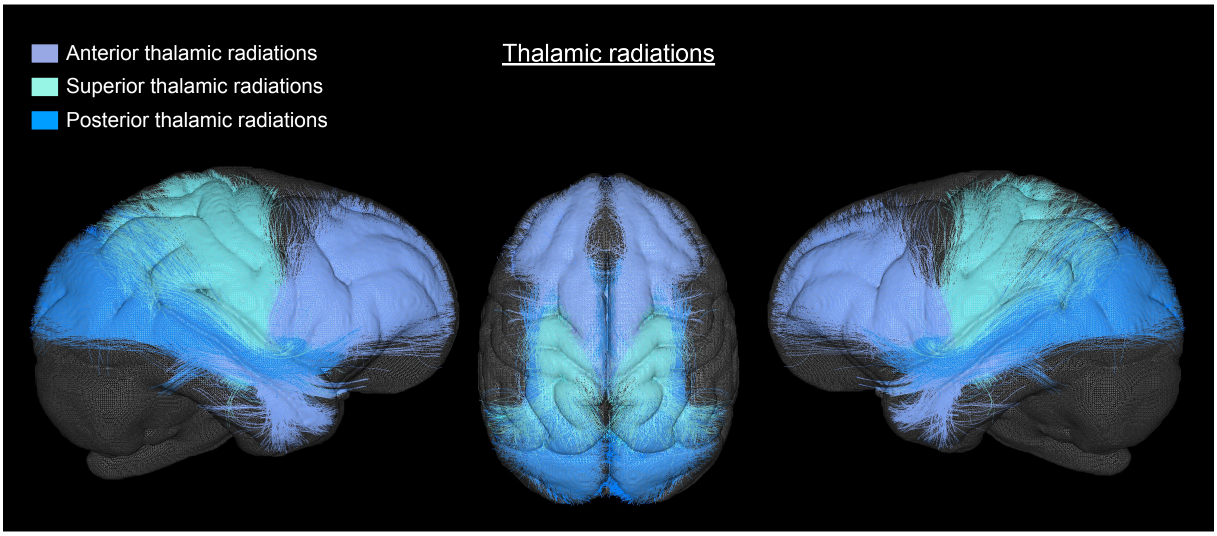


*Figure 5: The Thalamic radiations. Sagital left and right, and superior view of the thalamic radiations on a 3D mesh of the Juna template brain. Ant : Anterior, Sup : Superior, post : Posterior.*

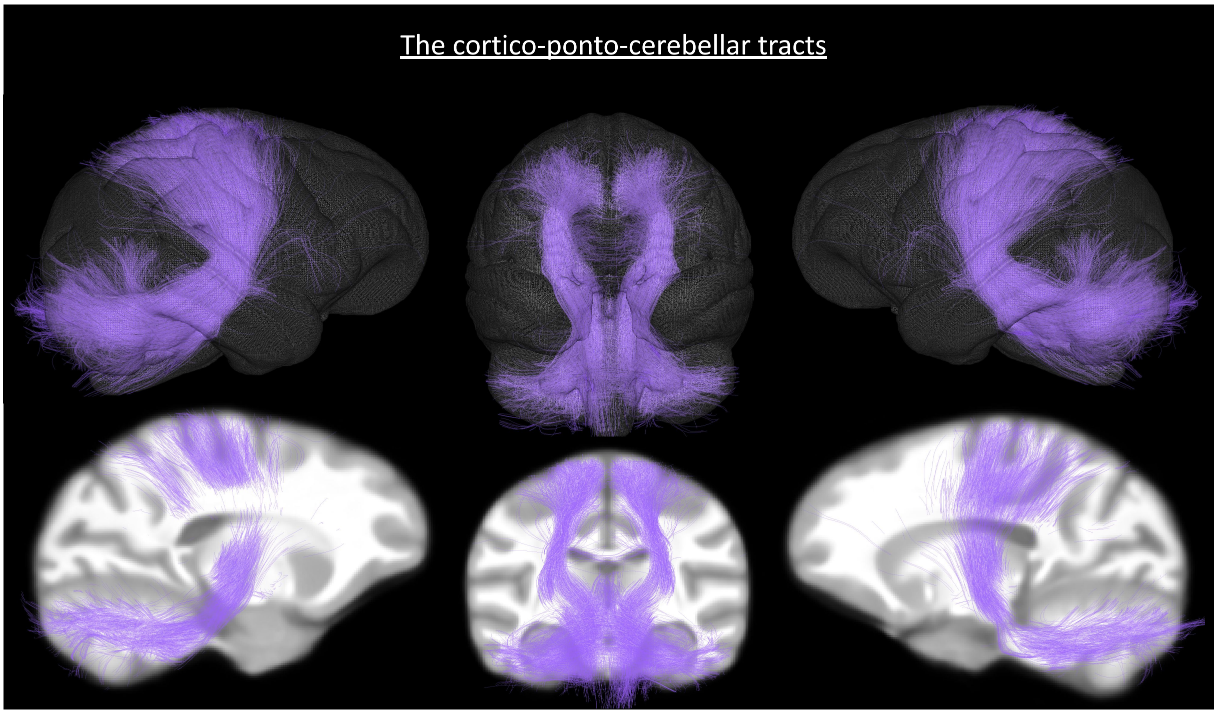


*Figure 6: The cortico-ponto-cerebellar tracts. (Top) tracts superimposed on a 3D mesh of the Juna.Chimp template; (Bottom) tracts superimposed on the T1-weighted anatomical image of the Juna.Chimp template. left sagittal, coronal, and right sagittal views.*

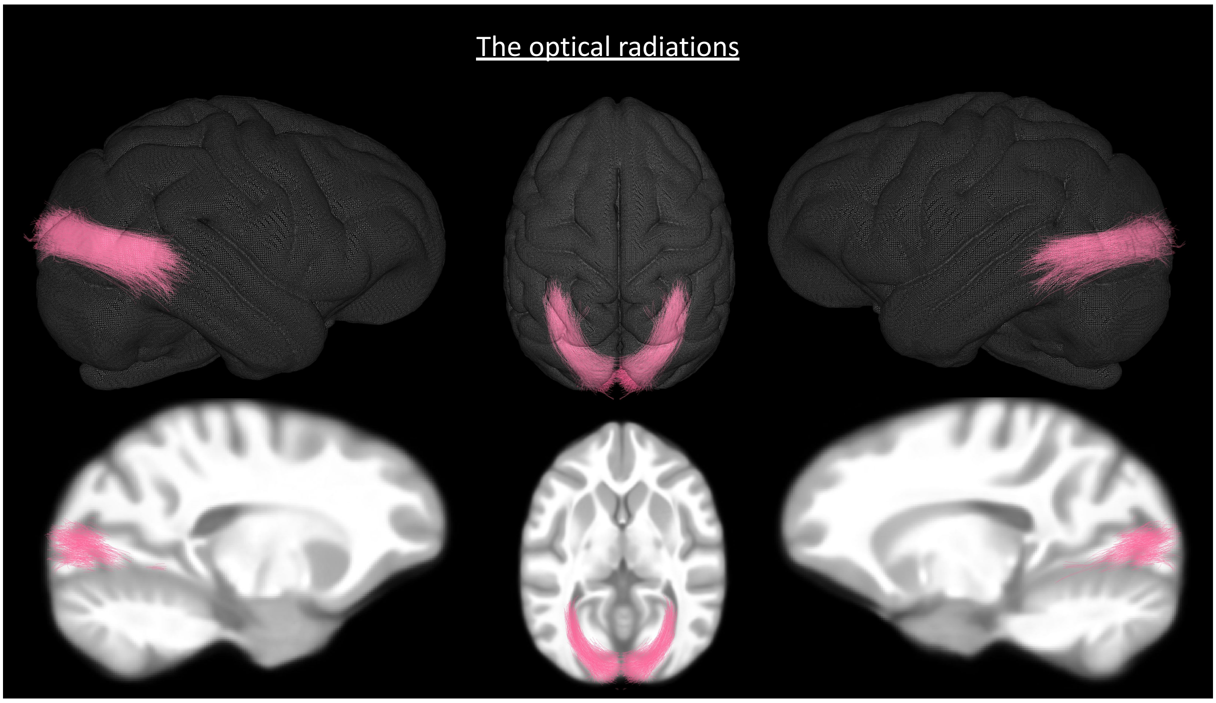


*Figure 7: The optical radiations. (Top) tracts superimposed on a 3D mesh of the Juna.Chimp template; (Bottom) tracts superimposed on the T1-weighted anatomical image of the Juna.Chimp template. left sagittal, coronal, and right sagittal views.*

Among the **association fibers** reconstructed in our deep white matter atlas are pairs of contralateral white matter bundles including : the inferior fronto-occipital fasciculi (IFOF), see figure 8, the inferior longitudinal fasciculi (ILF), see figure 9, the middle longitudinal fasciculi (MLF), see figure 10, the ventral visual streams (VVS), see figure 11, the arcuate fasciculi (AF), see figure 12, the frontal aslant tracts (FAT), see figure 13, the uncinate fasciculi (Unc), see figure 14, the cingulum (CG) (including ventral CGv and dorsal CGd), see figure 15 and the fornix (FX), see figure 16.

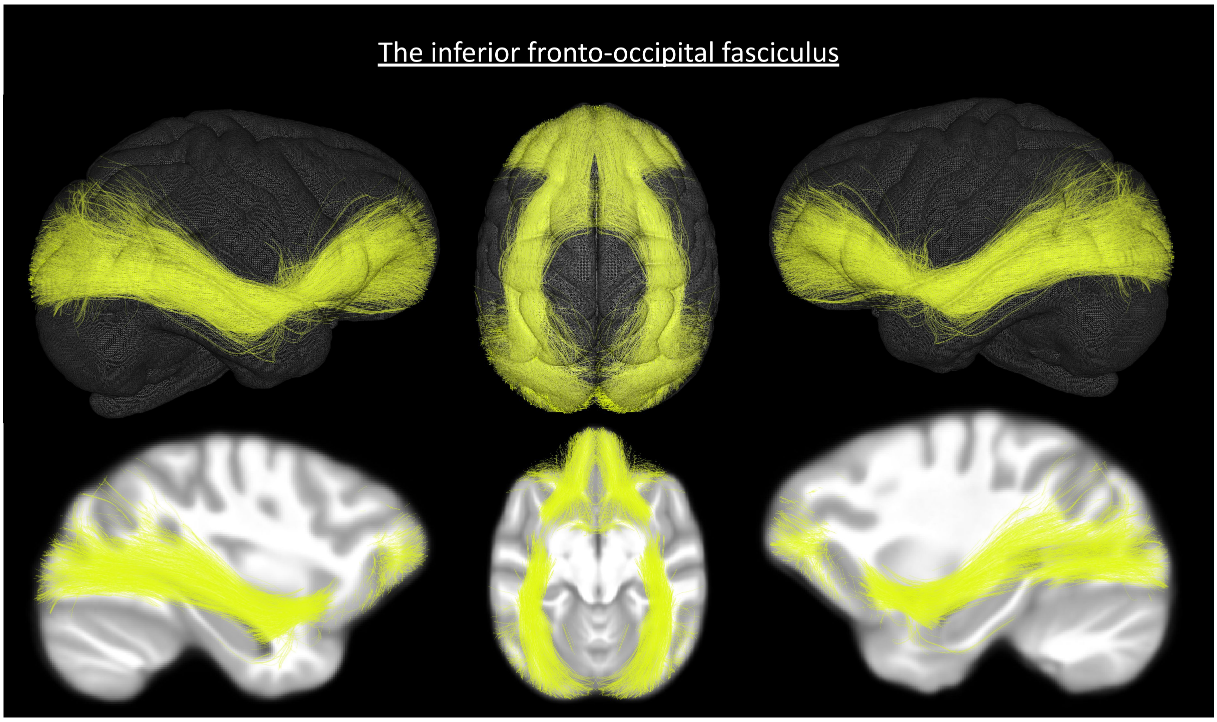


*Figure 8: The inferior fronto-occipital fasciculus. (Top) tracts superimposed on a 3D mesh of the Juna.Chimp template; (Bottom) tracts superimposed on the T1-weighted anatomical image of the Juna.Chimp template. left sagittal, coronal, and right sagittal views.*

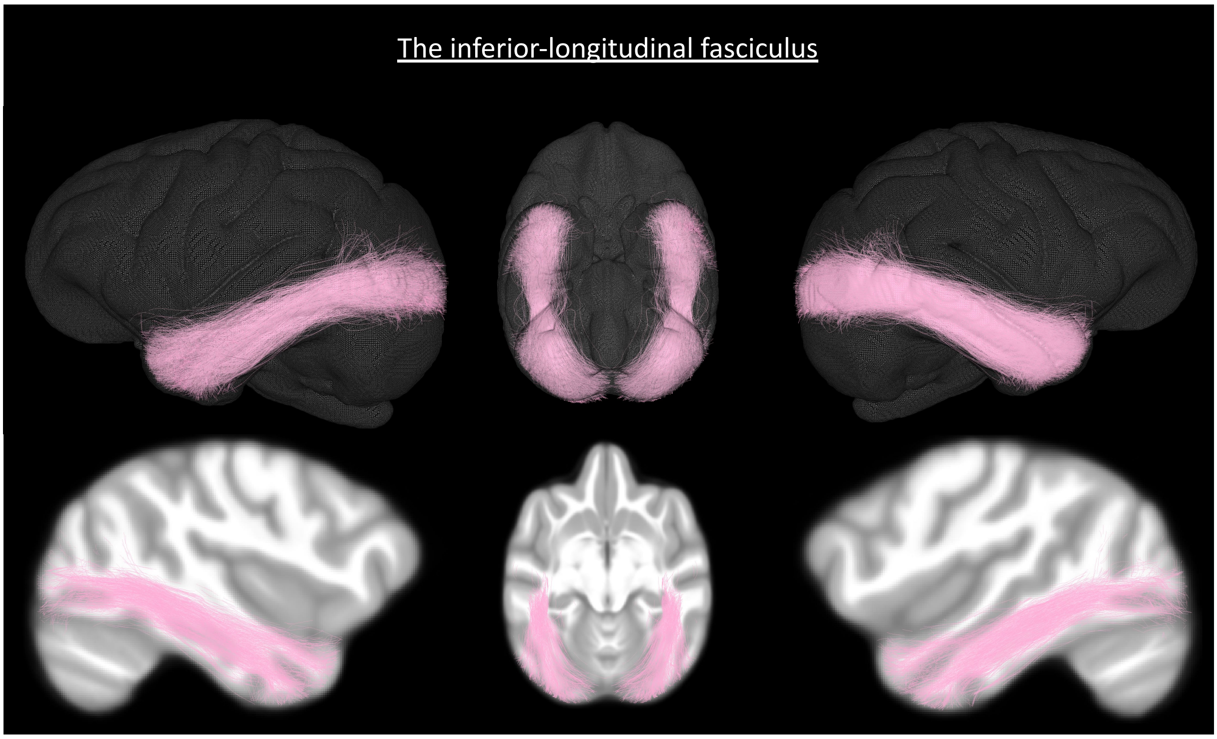


*Figure 9: The inferior longitudinal fasciculus. (Top) tracts superimposed on a 3D mesh of the Juna.Chimp template; (Bottom) tracts superimposed on the T1-weighted anatomical image of the Juna.Chimp template. left sagittal, coronal, and right sagittal views.*

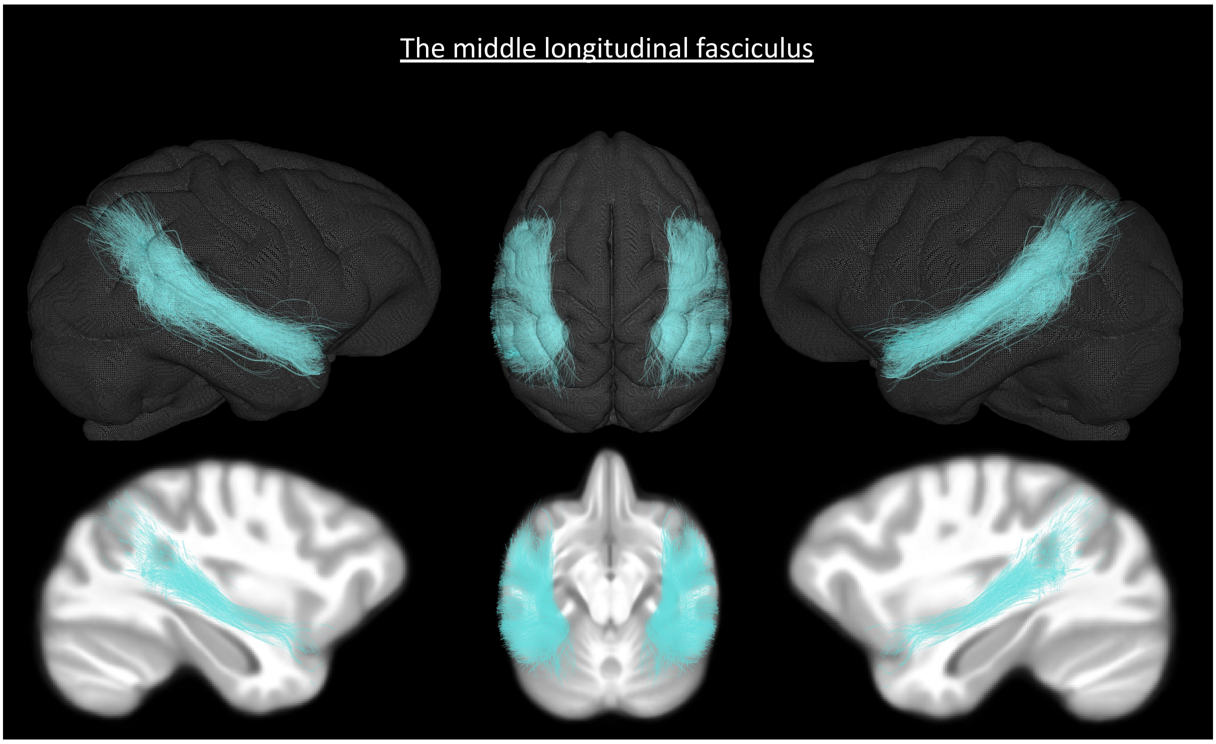


*Figure 10: The middle longitudinal fasciculus. (Top) tracts superimposed on a 3D mesh of the Juna.Chimp template; (Bottom) tracts superimposed on the T1-weighted anatomical image of the Juna.Chimp template. left sagittal, coronal, and right sagittal views.*

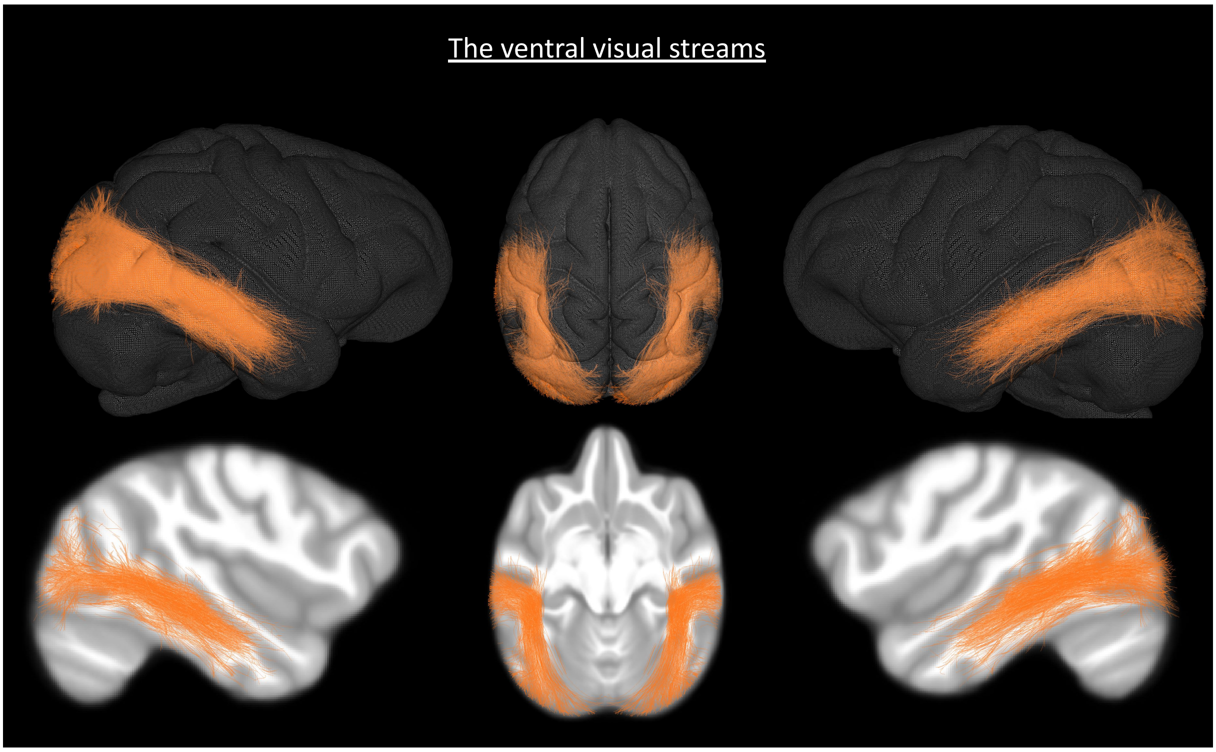


*Figure 11: The ventral visual streams. (Top) tracts superimposed on a 3D mesh of the Juna.Chimp template; (Bottom) tracts superimposed on the T1-weighted anatomical image of the Juna.Chimp template. left sagittal, coronal, and right sagittal views.*

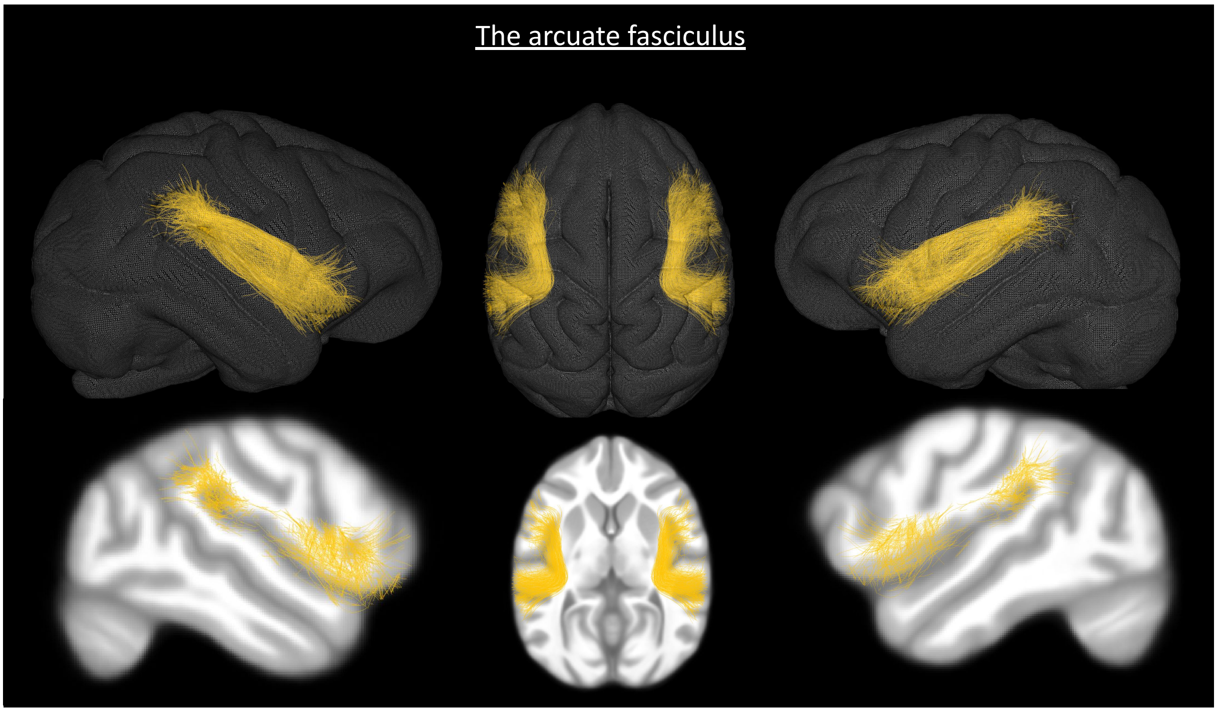


*Figure 12: The arcuate fasciculus. (Top) tracts superimposed on a 3D mesh of the Juna.Chimp template; (Bottom) tracts superimposed on the T1-weighted anatomical image of the Juna.Chimp template. left sagittal, coronal, and right sagittal views.*

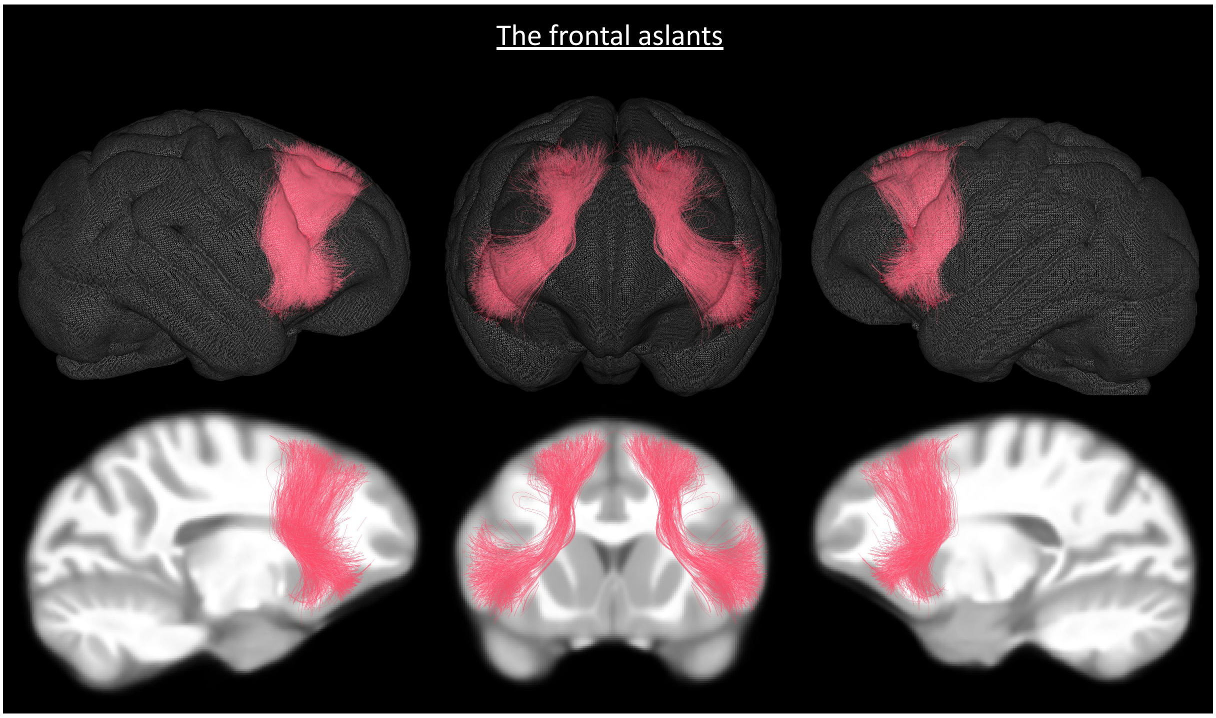


*Figure 13: The frontal aslants. (Top) tracts superimposed on a 3D mesh of the Juna.Chimp template; (Bottom) tracts superimposed on the T1-weighted anatomical image of the Juna.Chimp template. left sagittal, coronal, and right sagittal views.*

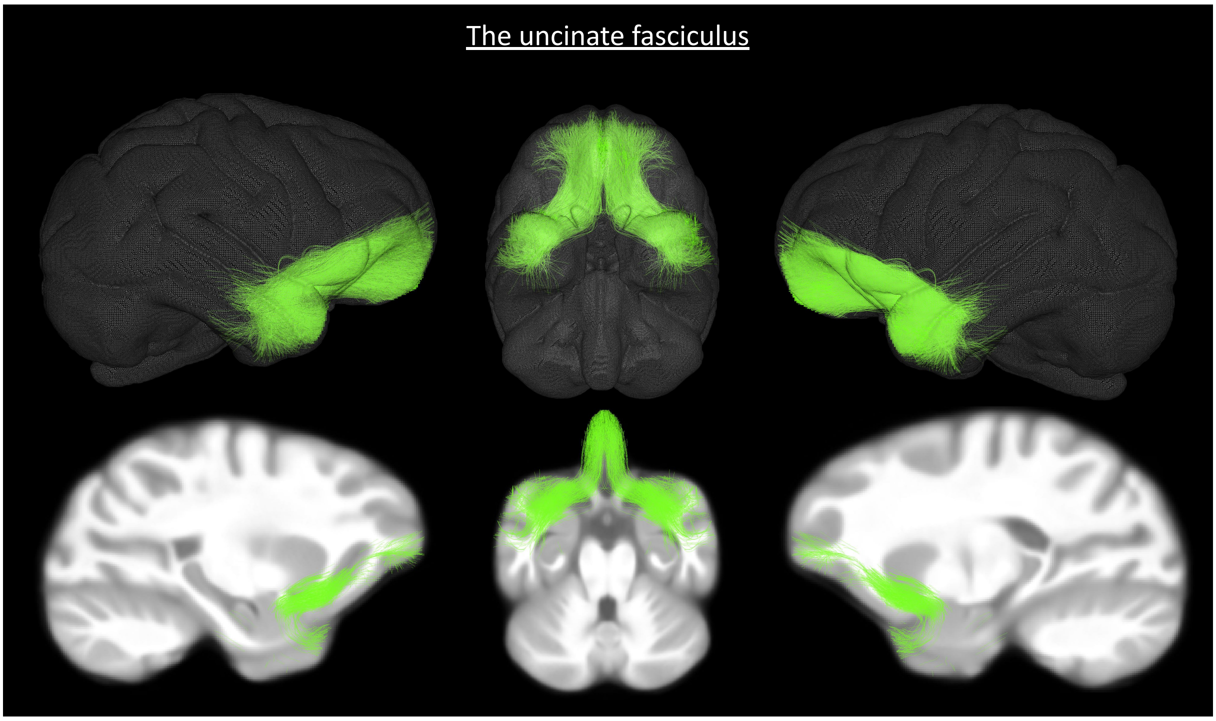


*Figure 14: The uncinate fasciculus. (Top) tracts superimposed on a 3D mesh of the Juna.Chimp template; (Bottom) tracts superimposed on the T1-weighted anatomical image of the Juna.Chimp template. left sagittal, coronal, and right sagittal views.*

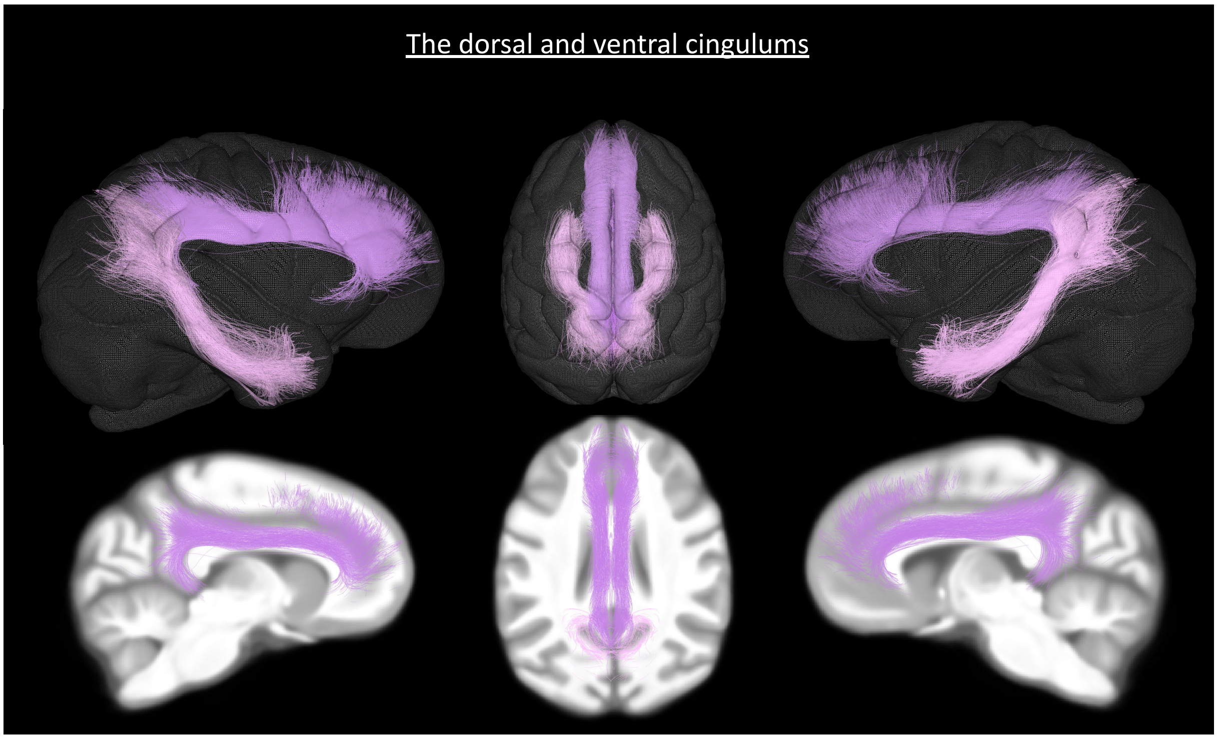


*Figure 15: The ventral and dorsal cingulums. (Top) tracts superimposed on a 3D mesh of the Juna.Chimp template; (Bottom) tracts superimposed on the T1-weighted anatomical image of the Juna.Chimp template. left sagittal, coronal, and right sagittal views.*

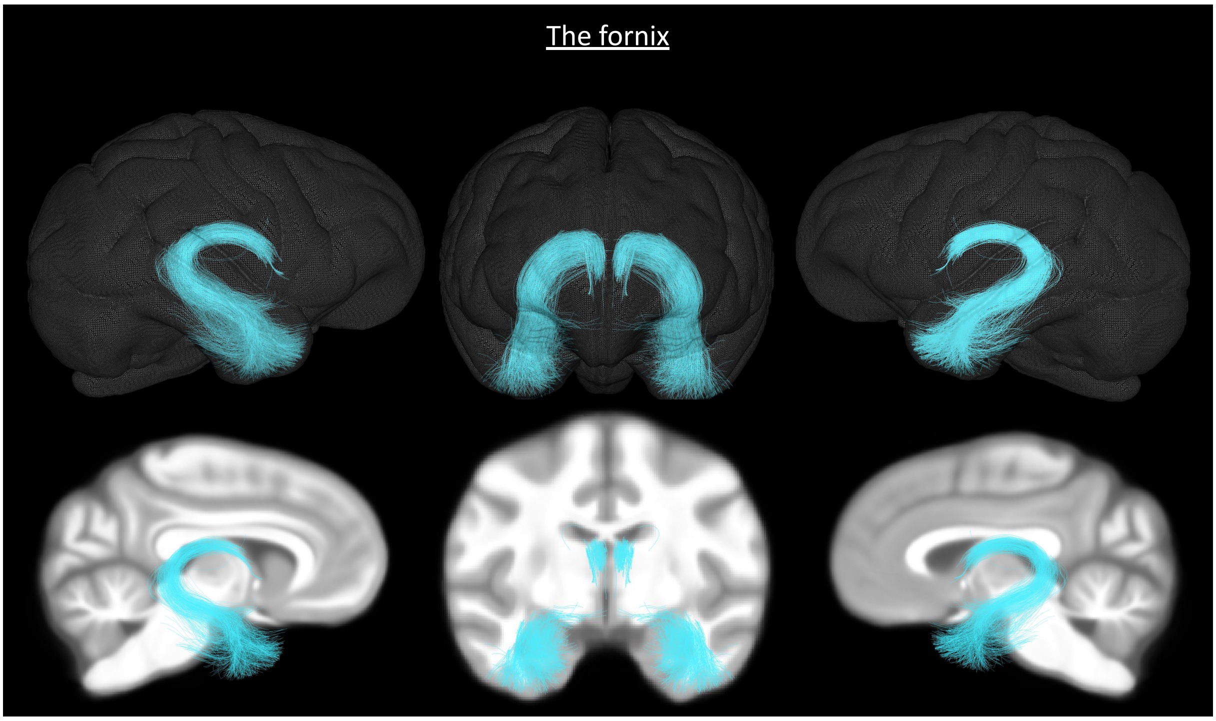


*Figure 16: The fornix. (Top) tracts superimposed on a 3D mesh of the Juna.Chimp template; (Bottom) tracts superimposed on the T1-weighted anatomical image of the Juna.Chimp template. left sagittal, coronal, and right sagittal views.*

**Ponto-cerebellar fibers** could also be identified including : the cortico-ponto-cerebellar tracts, described above, the parallel fibers (see figure 17) and the hypothalamic/subthalamic fibers (see figure 18).

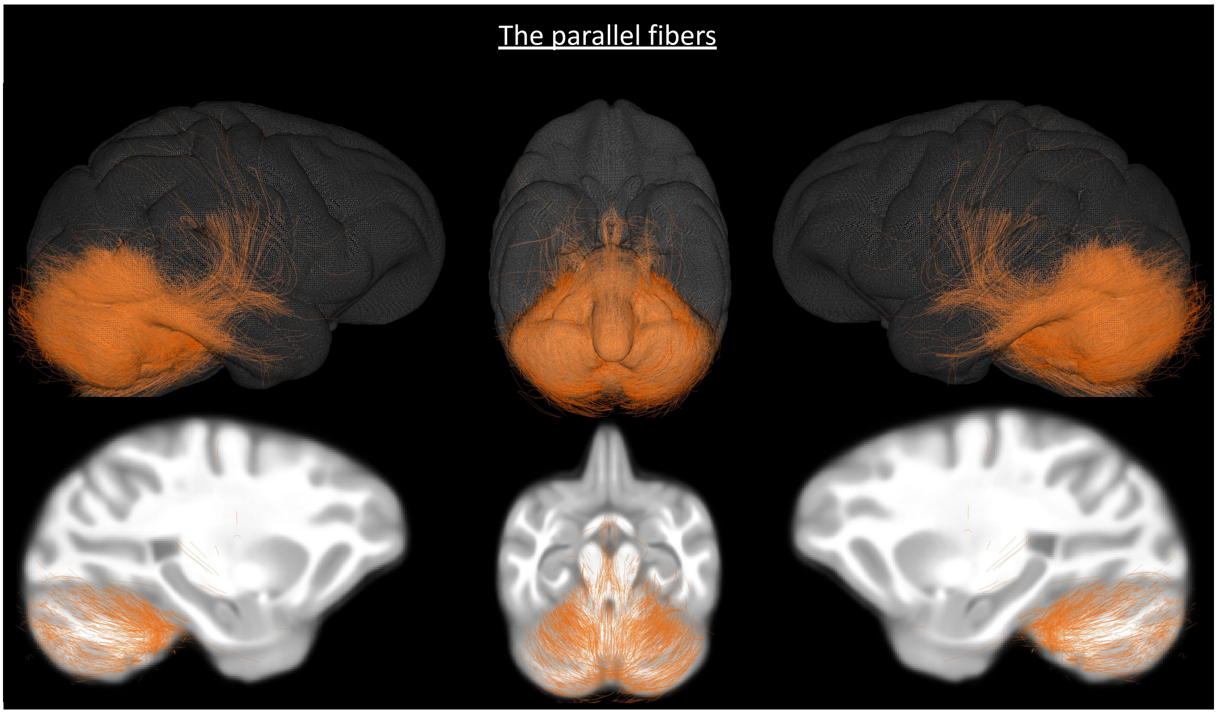


*Figure 17: The parallel fibers. (Top) tracts superimposed on a 3D mesh of the Juna.Chimp template; (Bottom) tracts superimposed on the T1-weighted anatomical image of the Juna.Chimp template. left sagittal, coronal, and right sagittal views.*

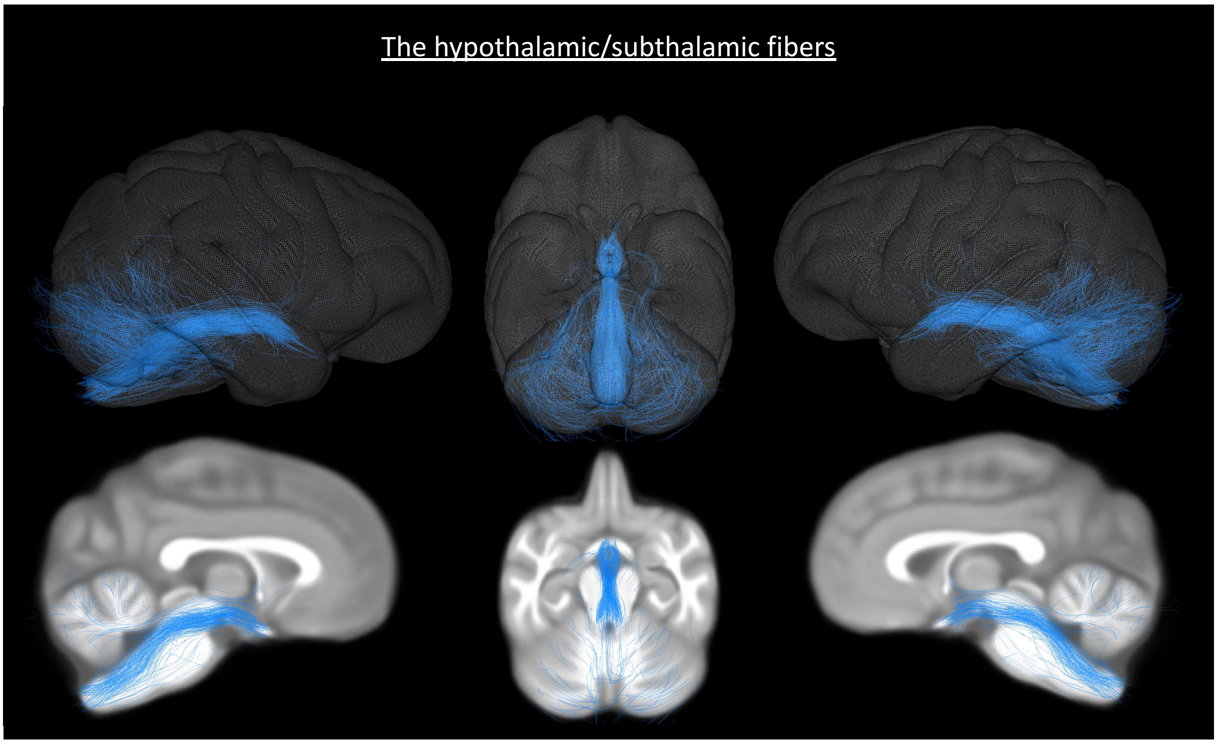


*Figure 18: The hypothalamic/subthalamic fibers. (Top) tracts superimposed on a 3D mesh of the Juna.Chimp template; (Bottom) tracts superimposed on the T1-weighted anatomical image of the Juna.Chimp template. left sagittal, coronal, and right sagittal views.*
